# An analytical pipeline for DNA Methylation Array Biomarker Studies

**DOI:** 10.1101/2021.07.14.452293

**Authors:** Jennifer Lu, Darren Korbie, Matt Trau

## Abstract

DNA methylation is one of the most commonly studied epigenetic biomarkers, due to its role in disease and development. The Illumina Infinium methylation arrays still remains the most common method to interrogate methylation across the human genome, due to its capabilities of screening over 480, 000 loci simultaneously. As such, initiatives such as The Cancer Genome Atlas (TCGA) have utilized this technology to examine the methylation profile of over 20,000 cancer samples. There is a growing body of methods for pre-processing, normalisation and analysis of array-based DNA methylation data. However, the shape and sampling distribution of probe-wise methylation that could influence the way data should be examined was rarely discussed. Therefore, this article introduces a pipeline that predicts the shape and distribution of normalised methylation patterns prior to selection of the most optimal inferential statistics screen for differential methylation. Additionally, we put forward an alternative pipeline, which employed feature selection, and demonstrate its ability to select for biomarkers with outstanding differences in methylation, which does not require the predetermination of the shape or distribution of the data of interest.

**Availability:** The Distribution test and the feature selection pipelines are available for download at: https://github.com/uqjlu8/DistributionTest

## Introduction

In most mammalian cells, DNA methylation is an epigenetic medication that occurs when a methyl group is added to position 5 of a cytosine within a CpG dinucleotide. This epigenetic mechanism acts as an important regulator of gene expression and maintenance of normal cell development [1]. In cancer, DNA experiences a global loss of methylation (hypomethylation), while selective hypermethylation occurs at regulatory regions (i.e. promotor-associated regions) leading to the silencing of key genes which are necessary for preventing the onset of different cancers [2]–[4]. Cancer cells use this reversible epigenetic programming mechanism to alter various signalling pathways to ensure their own survival, while consuming other tissues [5]. Therefore, DNA methylation have played a vital role in the development of novel biomarkers for the detection and prognosis of different cancers [6]–[8].

Increased interest in DNA methylation and its association with disease and cancer have resulted in the development of different assays to detect the change in pattern both at an epigenome-wide and locus-specific scale [9], [10]. The *Infinium HumanMethylation450* (HM450) BeadChip is a high-density microarray for quantifying the methylation level of over 450, 000 CpG sites within the human genome. The *Infinium* chemistry enables the reliable measurement of methylation statues to a single base resolution, and covers 99% of RefSeq genes with multiple sites in the annotated promoter (1500 base pairs (bp) or 200 bp upstream of transcription start sites), 5’-UTRs, first exons, gene bodies, and 3’-UTRs [11]. Due to the high-throughput capacity and the cost-effectiveness of this platform, initiatives such as The Cancer Genome Atlas (TCGA) and the International Cancer Genome Consortium (ICGC) have readily used the HM450 assay to explore the methylation profiles of different cancer samples [12], [13].

This has prompted the development of a variety of software packages to process and interpret the results, as well as quality control, interrogation, background correction and normalisation of processed results (e.g. *limma, minfi, missMethyl)* [14]–[16]. The methylation status is often measured using either log2 ratio of the intensities of methylated probes vs unmethylated probes (M-value), or beta (β) value, ranging from 0 – 1 (percentage of methylation), the latter of which has been used by different sources due to its intuitive biological interpretation. Generally, inferential statistical methods (e.g. t test or Mann-Whitney test) are often used to determine if there is a significant difference between the means of two sample groups. When a p-value is less than a given threshold (e.g. 0.05), the CpG site is considered to be differentially-methylated between the two samples, and referred to as a differentially-methylated CpG site.

Sampling distributions are important for inferential statistics given different types of data will require different types of analysis. The β value is preferentially used over the M value due to its easy biological interpretation. However, the M value will often be used for the statistical analysis, followed by visualization of results using the β value (as demonstrated by Maksimovic *et al* (2016) [17].

Through our initial survey of literature, there appeared to be a limitation of studies in this field, which considered the sampling distribution of individual CpG sites within a cohort of sample population from a statistical point of view prior to the selection of the most appropriate statistical method (**Table 1**).

**Table 1:**
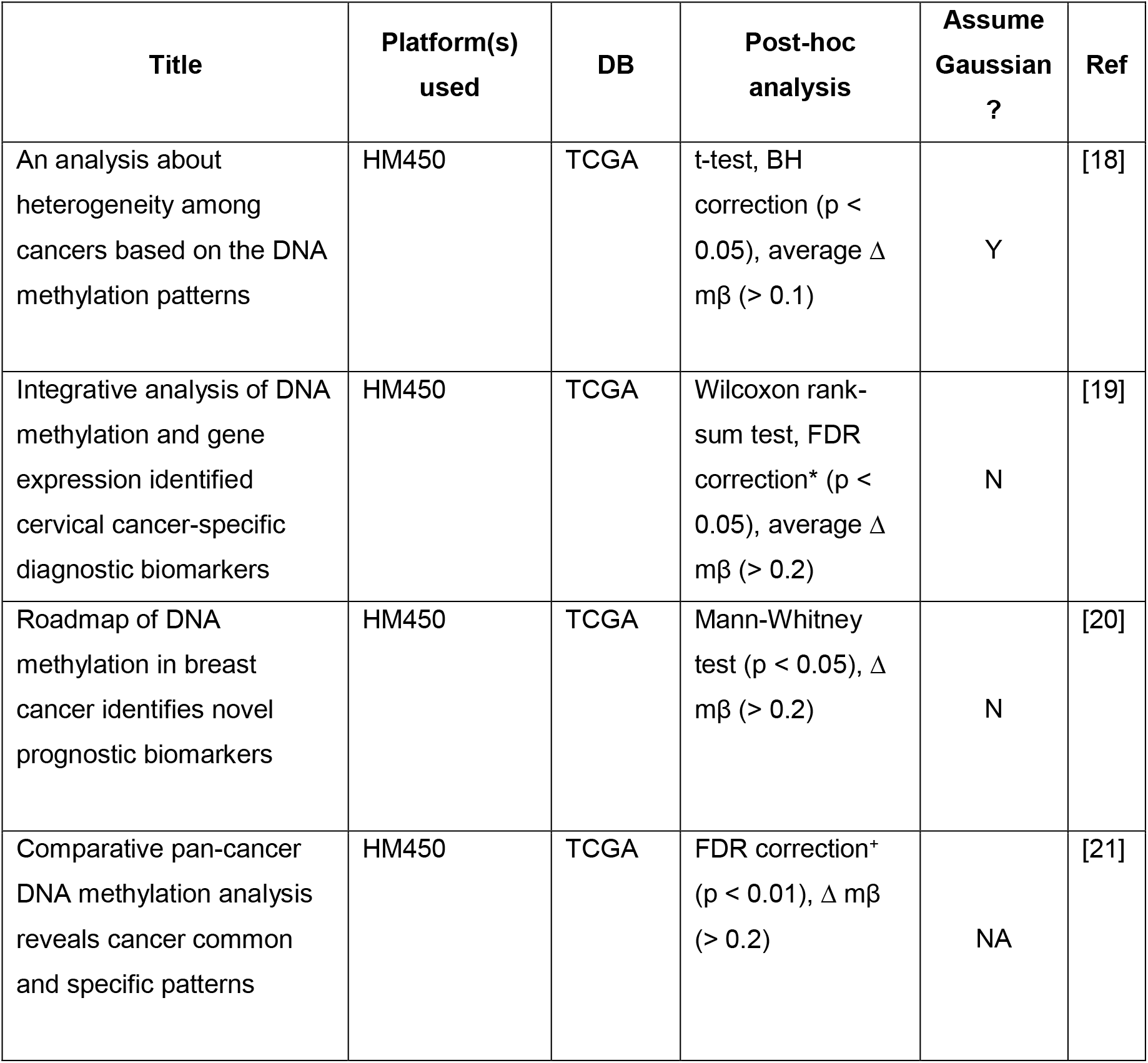
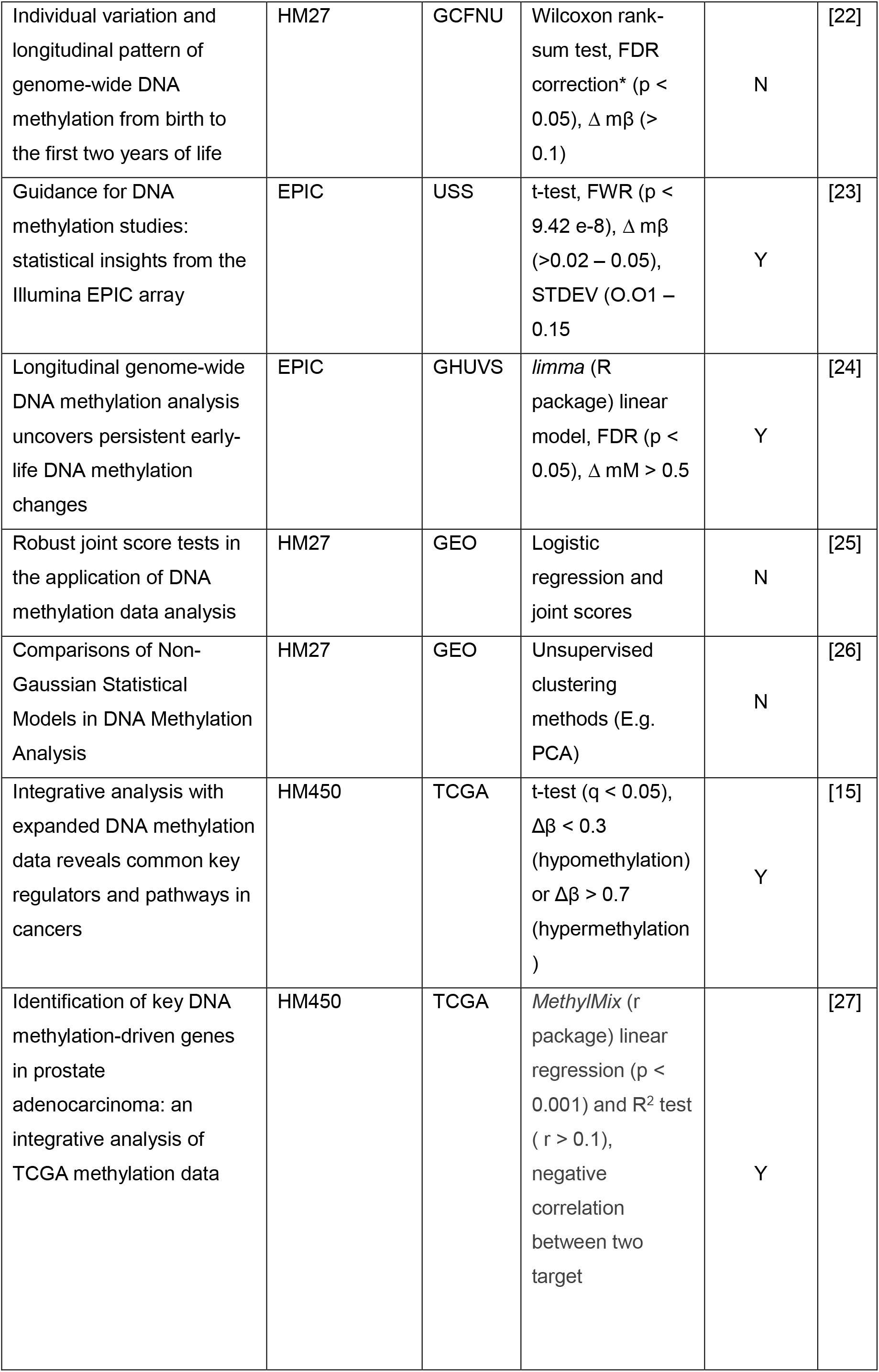
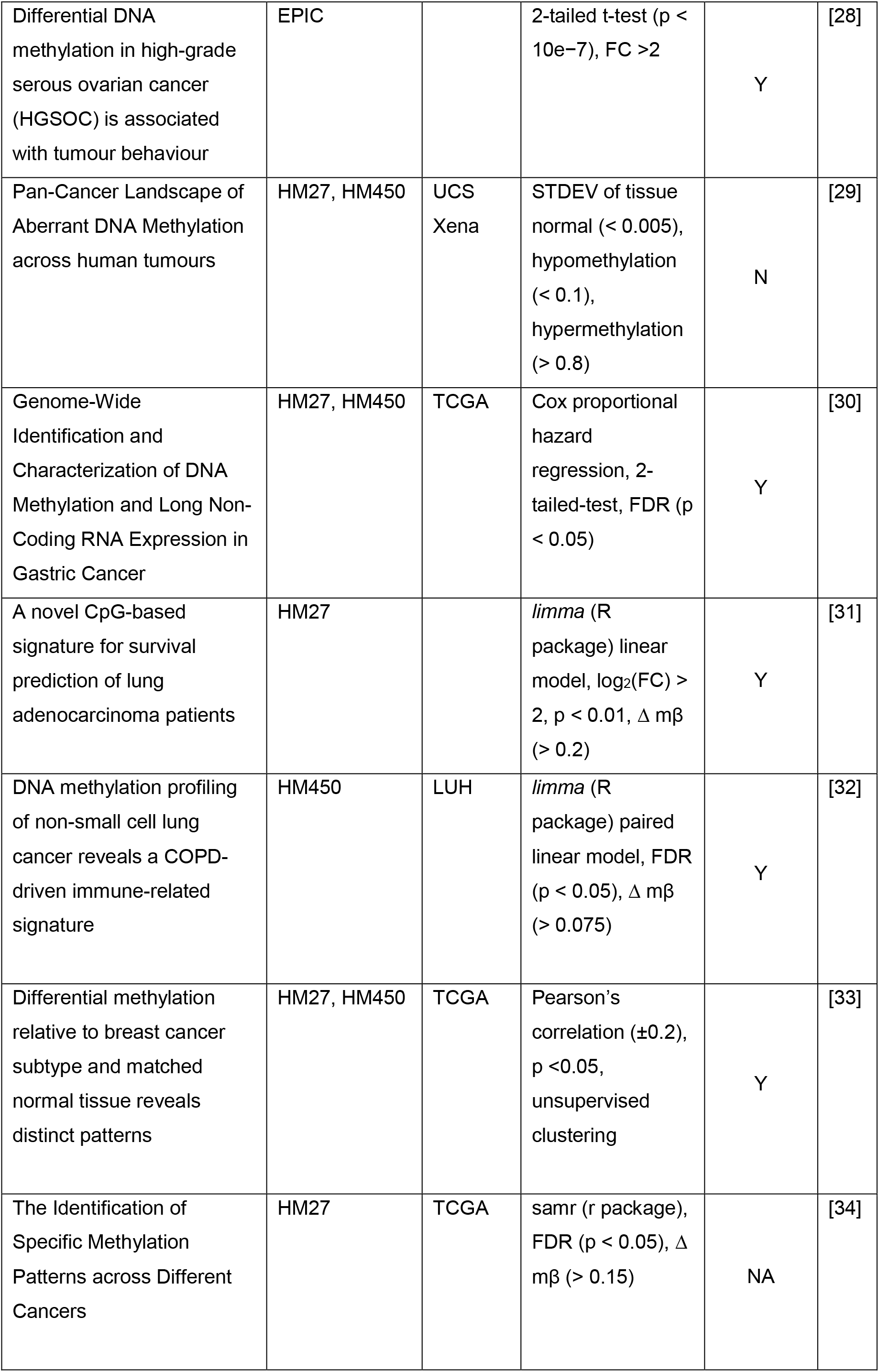
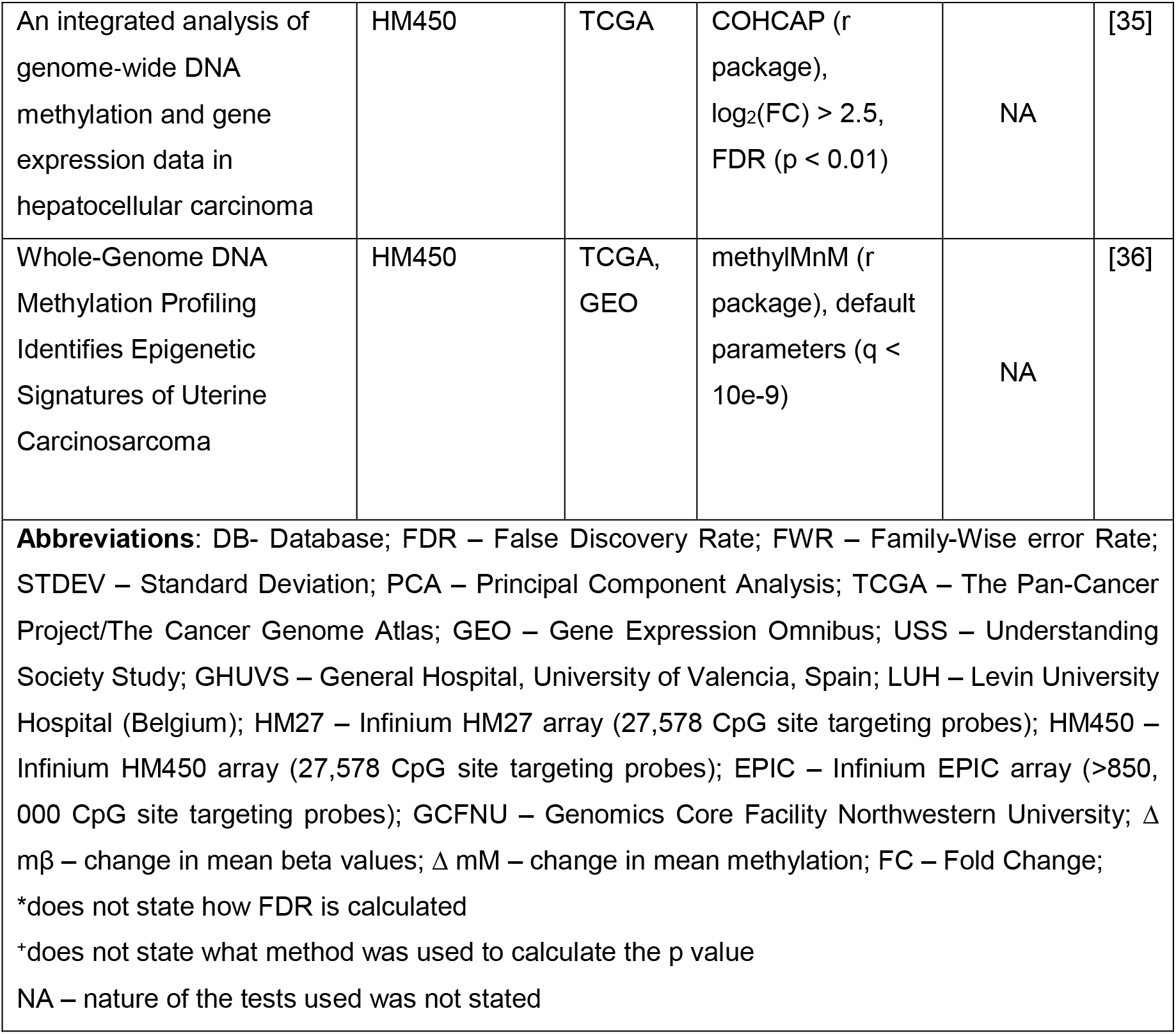
Summary of post-hoc analysis test(s) used by different epigenetic cancer studies that screened high-throughput microarray data for differentially-methylated regions.

Given epigenome-wide data generally followed a bimodal distribution, as demonstrated by Gallardo-Gomez *et al* (2018), whom analysed the methylation profile of pooled cell-free DNA (cfDNA) samples from colorectal cancer patients [37], the use of non-parametric tests in are appropriate as the latter does not make assumptions about the data. Similarly, bimodal distribution was observed in other microarray platforms (e.g. HM 27K, HM 450K array) and sequencing methods (e.g. bisulfite sequencing) when the results were analysed at an epigenome-wide scale [9], [16], [38].

Interestingly, when the distribution of probe (CpG)-wise methylation of samples within a single patient cohort (i.e. > 10 individuals) was visualised, a unimodal methylation distribution was observed [18], [35]. Given the probe-wise methylation of samples within a cohort has the potential to follow a Gaussian or non-Gaussian distribution, the determination of the spread and shape of the data is necessary to select the most optimal inferential statistical test to estimate the differences in methylation between sample populations. Due to the availability of a range of methylation analysis softwares (e.g. *minifi, watermelon, illumia* etc), which offers pipelines that preprocesses and analysis raw data into a readable format, the nature of the statistical method(s) used within each pipeline and the rationalization behind the tests used for each study is often limited [11], [39]–[41].

In most of the literature surveyed, differential methylation analysis was conducted by firstly performing a significance test to generate a p value of a cohort of samples across specific CpG sites (i.e. for the 27K array, up to 27,578 CpG sites surveyed). Due to the high volume of datasets analysed, a False Disovery Rate (FDR) was often used to lower the number of false positives encountered in the analysis. The FDR or α value generally falls between 0.01 – 0.05, where p ≤ α were considered to be statistically significant (**Table 1**). These biomarkers were then concluded to be differentially-methylated between specific cancer and matched (solid) tissue normal sample cohorts even though these difference were weakly present (approximately 10%) when the results were visualized using different methods (e.g. heatmaps) [18], [20], [21], [32]. Furthermore, given the same datasets from the TCGA, different significance testing methods (i.e. parametric and non-parametric) were used between the different studies, which implied that the overall shape and distribution of the methylation data may not be considered during analysis. However, past mathematical publications have suggested that different methods will give different p values based on a number of factors including: the sample size, shape and distribution of each dataset [42]–[44] and therefore may lead to a different level of significance and affect the overall interpretation of the differences in methylation.

Performing differential methylation analysis using the incorrect form of statistical methods may lead to the ‘false’ discovery of epigenetic biomarkers. This proved to be problematic in epigenetic biomarker discovery projects, where the main goal is to mine for CpG sites which presented a “clear” difference in methylation between a cancer and its matching tissue normal. Given most of these studies considered biomarkers with a difference in mean methylation of more than 10%, the use of significance testing as an initial post-hoc analysis may not be the most appropriate when screening for biomarkers with a distinct change in methylation between the tissue normal and the diseased cells [22], [23], [32].

Therefore, this study explored the use of different statistical methods to examine and screen for differentially-methylated CpG sites and highlighted the implications of using statistical methods that are sub-optimal for the given methylation data. From this, a methylation analysis pipeline was built in the python language which estimated the shape and distribution of each dataset prior to the selection of the ideal inferential statistical method on the given methylation samples. Additionally, we proposed the use of an alternative pipeline to screen for biomarkers with distinct methylation patterns between the tissue normal(s) and the cancerous samples. Based on feature selection, this pipeline was able to identify biomarkers with distinct methylation patterns of up to 12 different cancers and their matching tissue normal samples.

## Results

### Minimising the Level of False Positives by using the False Discovery Rate

The False Discovery Rate (FDR) refers to the expected ratio of type I errors (false positives) in hypothesis testing [45], [46]. In large-scale epigenome projects (e.g. TCGA project), where the number of comparisons (i.e. tests) for probe-wise methylation analysis may exceed 480,000, the False Discovery Rate (FDR) is often used to identify as many significant comparisons as possible while still maintaining a low false positive rate (i.e. the α value). In this study, the Benjamini-Hochberg (BH) method was used as it is not as conservative as traditional methods of correction (e.g. Bonferroni), which has been shown to lead to missed findings [45]–[48]. The α value was calculated from the cumulative p values of the comparisons between the (solid) tissue normal and the primary tumour samples of each cancer type using a custom python script (as summarised in **Figure 1**). With an average of 396, 065 comparisons (i.e. CpG targets) conducted across each cancer type, the α value was calculated to be 0.01 across fourteen cancer projects from the TCGA project.

**Figure 1:**
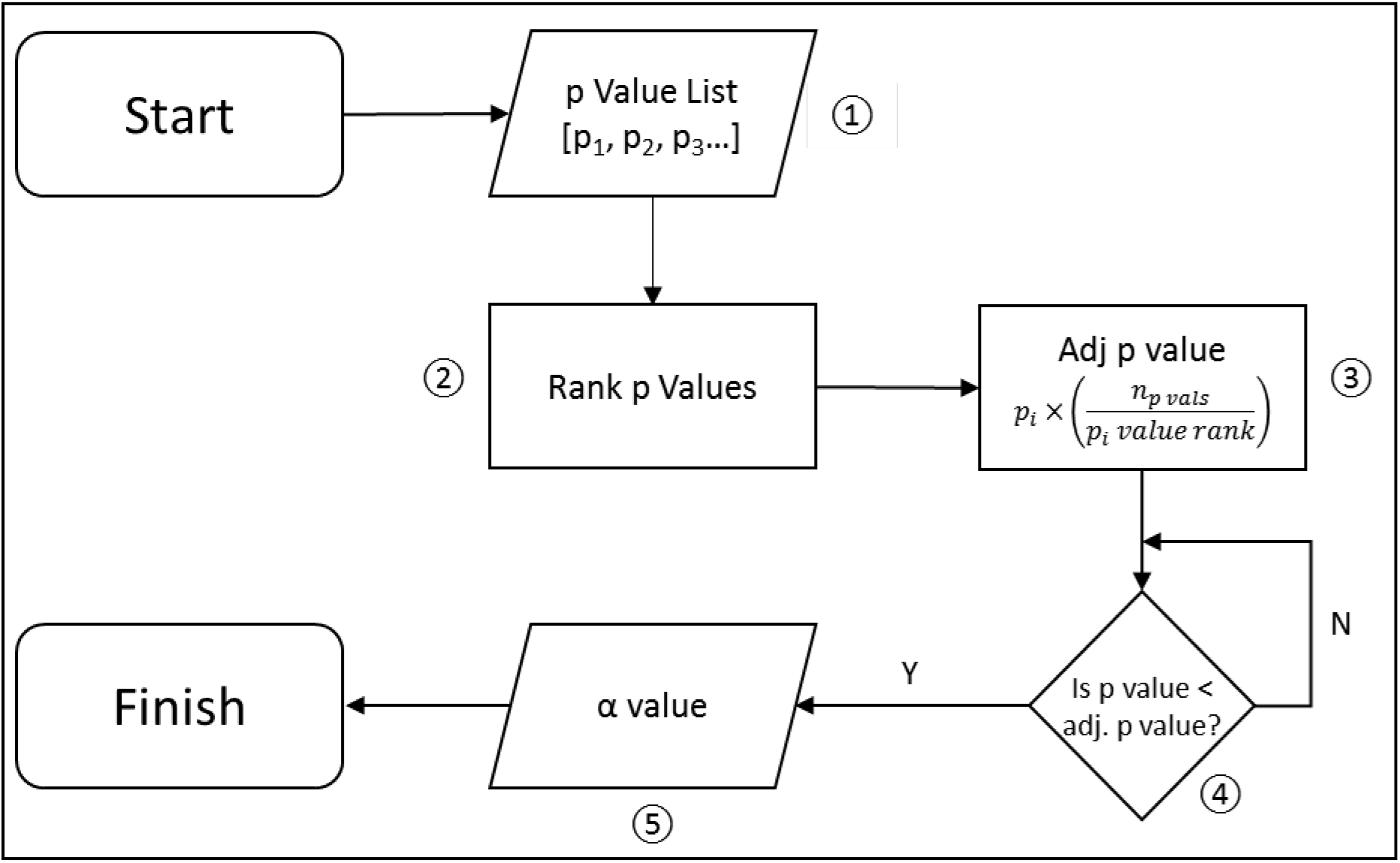
Calculating the False Discovery Rate using the Benjamini-Hochberg method. The Benjamini-Hochberg method controls for multiple hypothesis testing by controlling for the incorrect rejection of hypothesis. In the Benjamini-Hochberg method, the ① p values from all the results of a single inferential test (e.g. t-test) are collected and ② ranked from smallest to largest. ③ The Adjusted (Adj.) p value at each position (i) is calculated by multiplying the p value by the number of p values (n_p vals_) over the rank of the i^th^ p-value. ④ The p-value and adjusted p-value is observed ⑤ where α value is the adjusted p value where the p value < the adjusted p value.

### Prediction of methylation distribution using the “Distribution test”

The *Distribution test* was written in the python language to predict the distribution and shape of input methylation data prior to the selection of the optimal inferential statistical test (e.g. t-test, Mann-Whitney test) to screen for differential methylation between two or more sample groups. The distribution of each dataset was calculated using three commonly-used distribution tests: Shapiro-Wilk, D’Agostino and Anderson-Darling [49]–[51] (as summarised in **Figure 2**). Additionally, the symmetry and the combined sizes of the two tails of the distribution was calculated using the skewness and kurtosis tests respectively (as summarised in **Figure 3**) [52].

**Figure 2:**
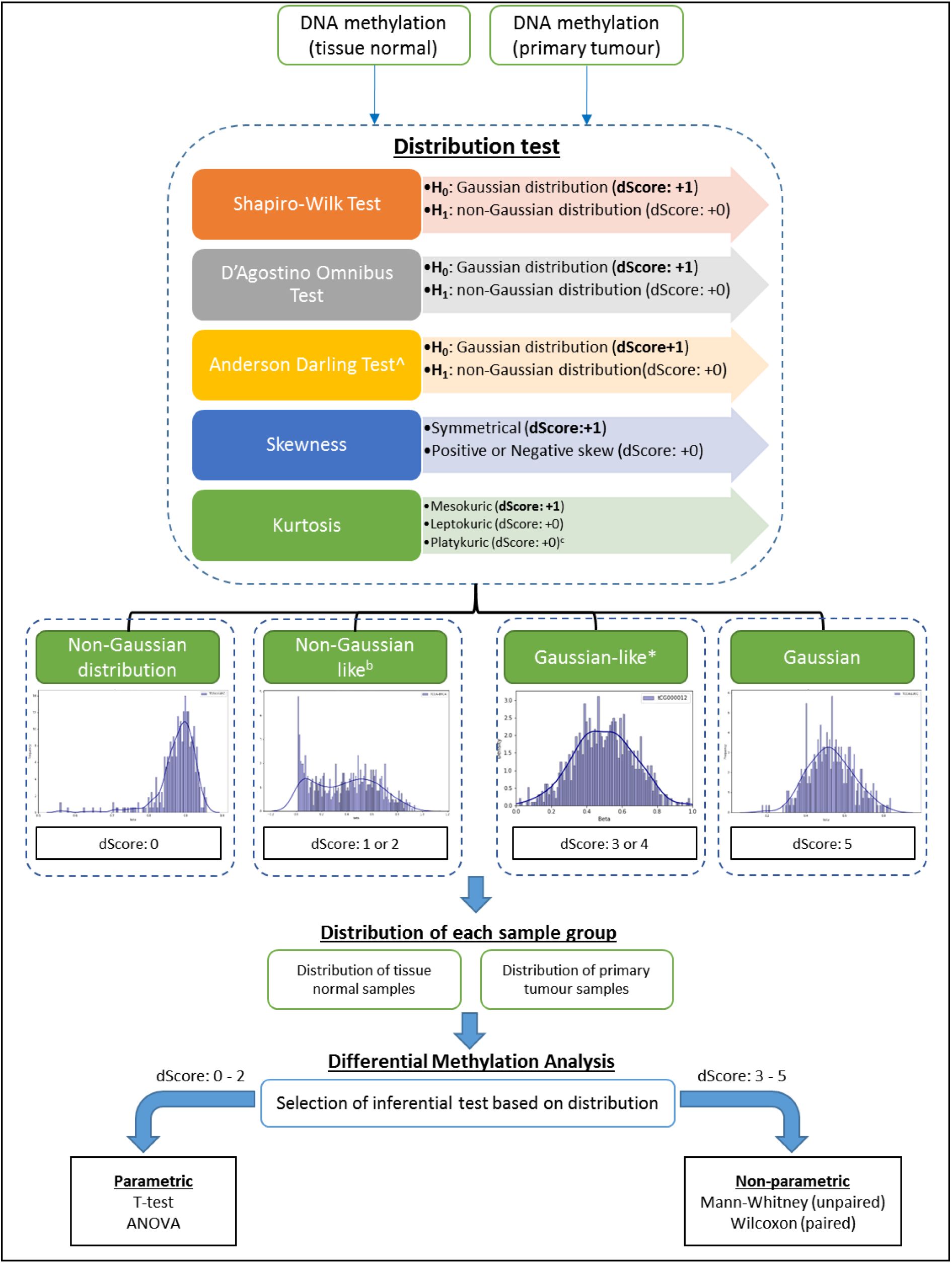
Workflow of the Distribution Test pipeline. The DNA methylation pattern of the primary tumour and (solid) tissue normal of each cancer site was analysed separately through the “Distribution test” pipeline. The pipeline determines if each methylation dataset can be fitted into a “Gaussian” or “non-Gaussian” distribution from the results of the five tests used. Each test generates a binary number of 0 (non-Gaussian distribution) or 1 (Gaussian distribution) which when added together generates an overall distribution score (dScore) out of 5. A score of 0 suggests the data followed a non-Gaussian distribution, while a dScore of 5 indicates the data followed a Gaussian distribution. Additionally, a dataset can also be determined to follow a “non-Gaussian-like” or “Gaussian-like” distribution if the dScores are 1-2 or 3-4 respectively. Inferential (hypothesis) testing to determine the difference in methylation will be selected based on distribution of each sample group. Depending on the outcome of the distribution test, Differential methylation analysis using inferential testing is performed. ^ The Anderson-Darling test compares a test statistic to a critical value to determine the distribution, where H_0_: statistic < critical value = Gaussian distribution; H_1_: statistic > critical value = non-Gaussian distribution *where the majority of the distribution tests used suggests a “Gaussian distribution ^b^ where the majority of the distribution tests used suggests a “non-Gaussian” distribution ^c^ Platykuric shape was not applicable in methylation distribution datasets

**Figure 3:**
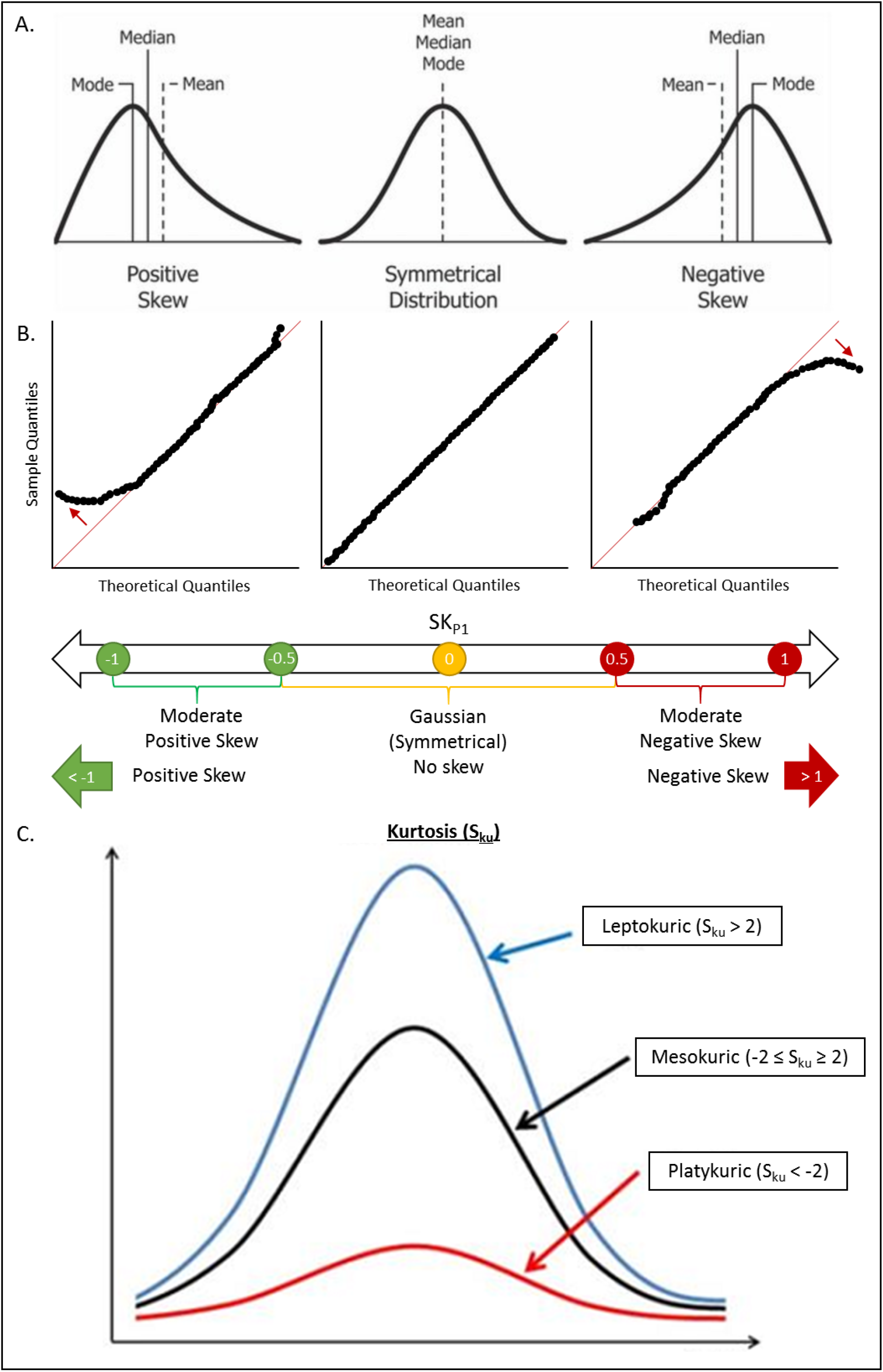
Measures of Skewness and Kurtosis. Skewness defines the degree of distortion from the symmetrical bell curve or the Gaussian distribution. **A**. Distribution of dataset as shown a histogram - (**left**) positive skewness: when the tail of the right side of the distribution is longer or heavier. The median and mean will be more than the mode; (**centre**) Gaussian (symmetrical) distribution: when there is minimal to no skewness resulting in a bell shape; (**right**) negative skewness: when the tail of the left side of the distribution is longer or heavier. The mean and median will be greater than the mode of the distribution. **B**. Distribution of dataset as shown by Q-Q Plots – (**left**) positive skewness: deviation from the trendline (→) indicates its skewness to the left; (**centre**) Gaussian distribution: distribution along the trendline indicates the data followed a Gaussian distribution; (**right**) negative skewness: deviation from the trendline (→) indicates its skewness to the right.

Using a combination of these five tests (i.e. Shapiro-Wilk, D’Agostino Omnibus, Anderson-Darling, skewness and kurtosis) the shape and distribution of each methylation dataset was calculated by generating a “distribution” score (dScore). The dScore for each dataset is set between 0 – 5, where 0 suggests the data followed a non-Gaussian distribution, while a dScore of 5 suggests the data followed a Gaussian distribution (as shown in **Figure 2**). However, a dScore of 1-2 suggests the dataset followed a “non-Gaussian-like” distribution, while a dScore of 3-4 suggests the dataset followed a “Gaussian-like” distribution. Depending on the outcome of the distribution test, the pipeline will screen for changes in methylation using inferential statistics differently. If both the tissue normal and primary tumour sample groups were scored to be between 0 – 2, parametric testing will be used. However, if the distribution of both sample groups scored between 3 – 5, or one group scored 0-2 and the other scored 3-5, than non-parametric testing will be used.

The Shapiro-Wilk and D’Agostino Omnibus test uses the null hypothesis (H_0_) to determine if a dataset is derived from a Gaussian distribution (i.e. bell-shaped curve), where if the p ≤ α (i.e. H0 is rejected), a non-Gaussian distribution was observed. However, if the p > α (accept H0 is accepted), then the dataset most likely followed a Gaussian distribution. Given the large number of probes for each dataset (396, 065), an α value of 0.01 was determined for each of the 14 cancer projects examined in this study using the Benjamini-Hochberg method [46], [48] (**Figure 1**). Of the 396,065 probes covered by each tissue of origin, the number of probes that followed a Gaussian distribution and the number of probes that followed a non-Gaussian Distribution using the custom Distribution test was summarised in **Figure 4**. To better visualise and compare the density of each distribution, a colour scale was used, were a high number of probes are highlighted in green while lower number of probes are highlighted in red. From **Figure 4A**, it was shown that most CpG sites (probes) of primary tumours followed a non-Gaussian or non-Gaussian-like distribution, while the opposite was true for (solid) tissue normal samples from the matched tissue normal. Furthermore, split violin plots were used to visualise the spread of each distribution (i.e. Gaussian, Gaussian-like, non-Gaussian, and non-Gaussian-like) for both the tissue normal and primary tumour samples of each cancer (TCGA-BRCA shown in **Figure 4B**, other violin plots shown in **Supplementary Figures 1-13**). For the majority of projects visualised using violin plots, there is a change in density when looking at the same beta values of corresponding distributions.

**Figure 4:**
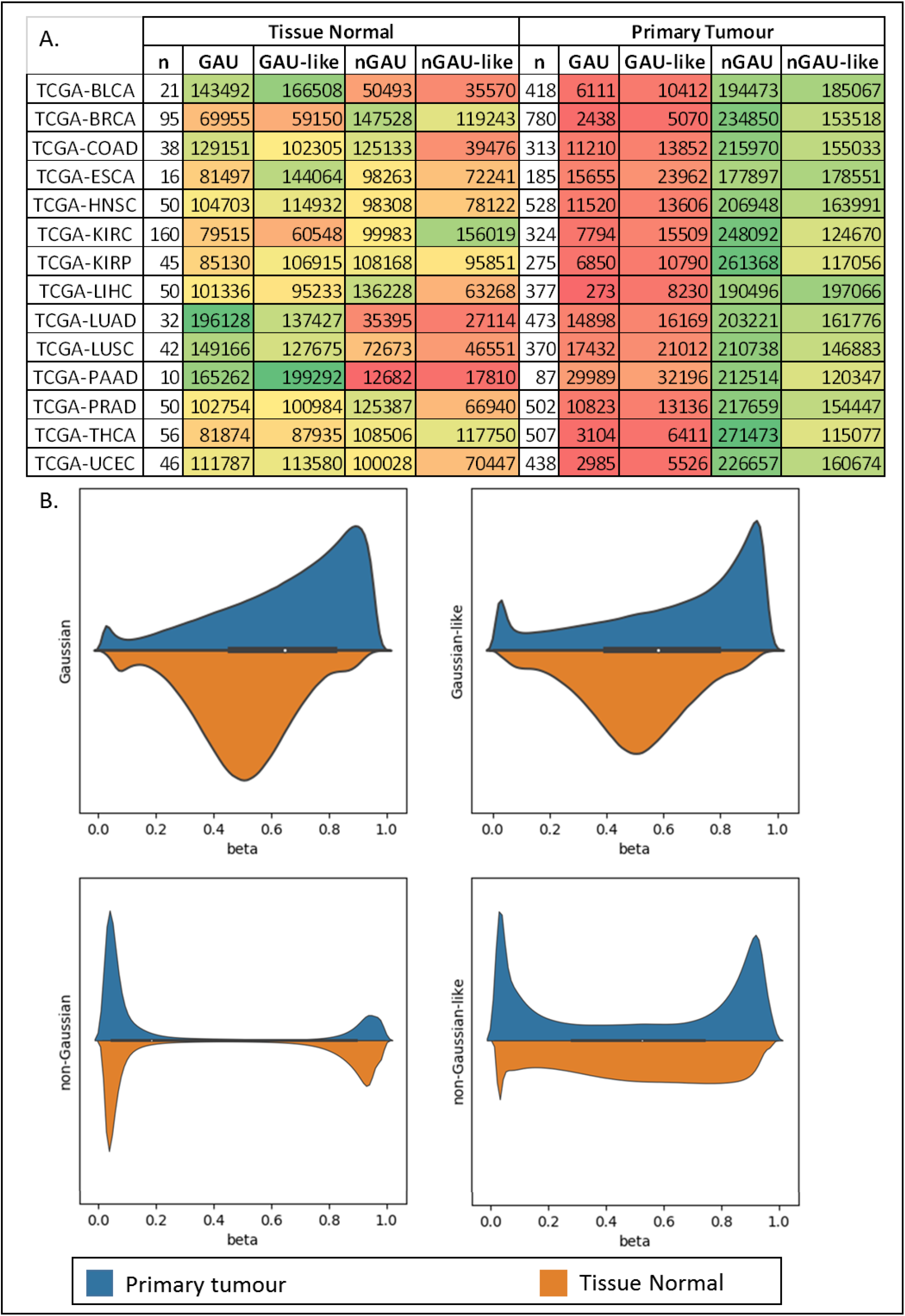
Summary of Distributions of TCGA-Dataset as determined by the custom Distribution test. **A.** Summary of the number of probes for matching tissue normal and primary tumour of each cancer which followed a specific distribution as determined using the Distribution test. By using a colour scale (red (low number) → green (high number), it was found that the majority of primary tumour samples followed a non-Gaussian (nGAU) or non-Gaussian-like distribution. In contrast, the matching (solid) tissue normal samples appeared to show an opposite trend in most cases. **B.** Split violin plots of primary tumour (orange) and tissue normal (blue) of the four different distributions () for TCGA-BRCA. Violin plots for remaining datasets are shown in **Supplementary Figure 1**. By comparing the distribution of the matching primary tumour and tissue normal for each distribution, it is clear there is a change in the number of probes at each beta level. *GAU – All Gaussian; GAU-like – Gaussian-like; nGAU – All non-Gaussian; nGAU-like – non-Gaussian-like. TCGA-BLCA – Bladder Urothelial Carcinoma; TCGA-BRCA – Breast Invasive Carcinoma; TCGA-COAD – Cholangiocarcinoma; TCGA-ESCA – Esophageal Carcinoma; TCGA-HNSC – Head and Neck Squamous Cell Carcinoma; TCGA-KIRC – Kidney Renal Clear Cell Carcinoma; TCGA-KIRP – Kidney Renal Papillary Cell Carcinoma; TCGA-LIHC – Liver Hepatocellular Carcinoma; TCGA-LUAD – Lung Adenocarcinoma; TCGA-LUSC – Lung Squamous Cell Carcinoma; TCGA-PAAD – Pancreatic Adenocarcinoma; TCGA-PRAD – Prostate Adenocarcinoma; TCGA-THCA – Thyroid Carcinoma; TCGA-UCEC – Uterine Corpus Endometrial Carcinoma.

Furthermore, since the Shapiro-Wilk, D’Agostino and Anderson-Darling test reported a binary answer (Gaussian, or non-Gaussian), Venn diagrams were used to visualise the number of probes that shared the same distributions without considering the shape (skewness) and the tail (kurtosis) of each dataset (**Figure 5**). From the Venn diagrams, most probes from primary tumour samples that followed a mutual non-Gaussian distribution when examined using the three normality tests. In contrast, samples from (solid) tissue normal that followed a mutual Gaussian distribution when visualised using a Venn diagram. Therefore, it was hypothesised that the number of samples used in each test may influence the final output of the distribution as well as overall shape. This was addressed below where the ability of the distribution test to predict the distribution and shape of each dataset was determined using simulated data.

**Figure 5:**
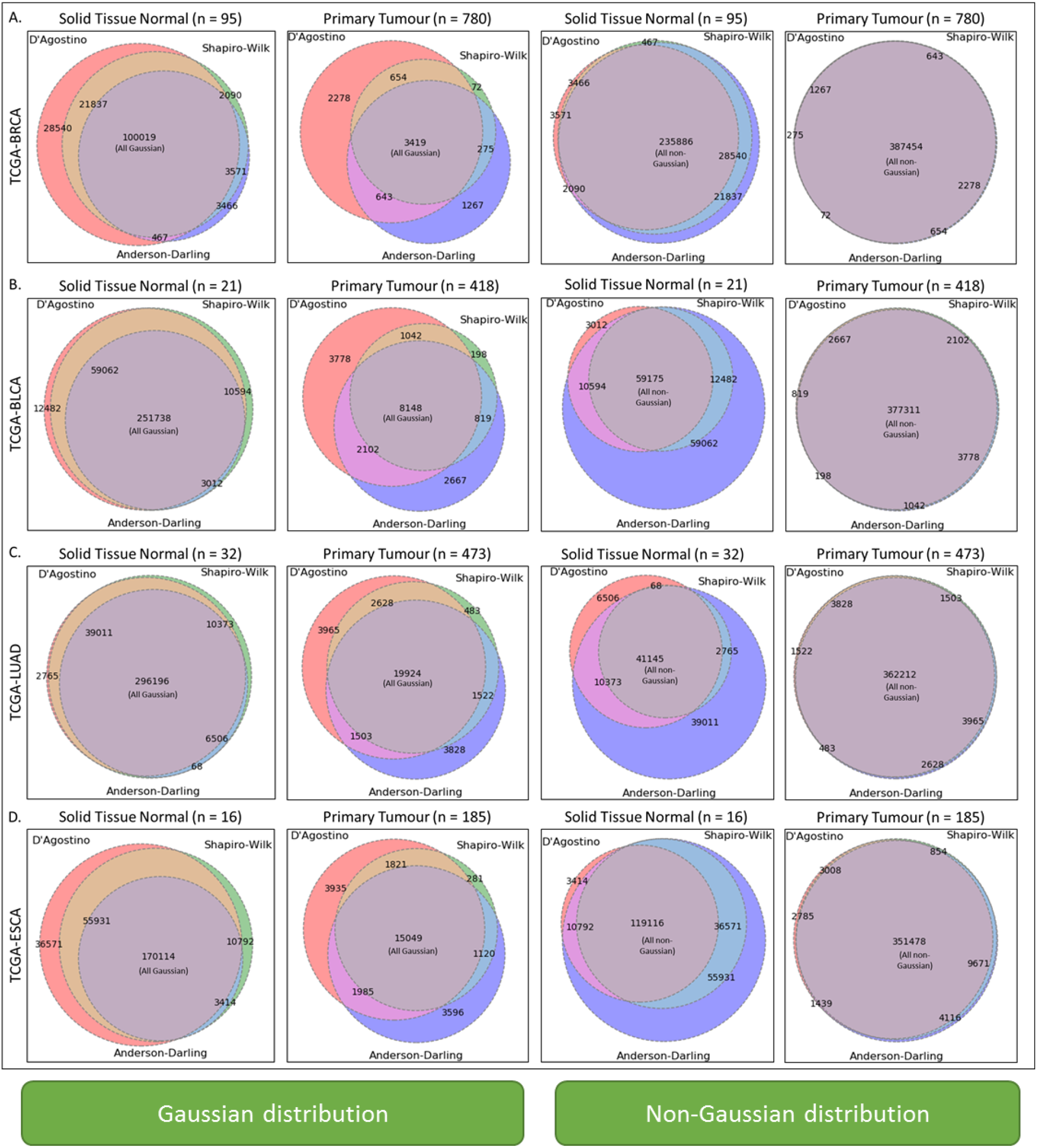
Summary of three distribution tests used by the Distribution test. Venn diagrams are used to show the intersection between three of the main tests used to determine the distribution in there of the tests (Shapiro-Wilk, D’Agostino, and Anderson-Darling) used in the Distribution test:. The number of probes which were predicted to follow a Gaussian distribution are shown in the two left columns, while the number of probes which followed a non-Gaussian distribution are shown in the two columns on the right. In this analysis, most CpG sites identified in Solid Tissue Normal samples are found to follow a Gaussian distribution, while most CpG sites from primary tumours are found to follow a non-Gaussian distribution.

### Data Simulation to validate the “Distribution test”

To determine if the tests in the “Distribution test” pipeline was correctly calling the distribution and shape of the dataset, simulated methylation (β values) data with a predefined shape and distribution was generated. This was conducted using a custom python script, with the number of samples in each dataset ranging from 100 to 1000 (Statistical summary results from Distribution test available upon request), and visual validation of representative simulated targets performed using both a histogram and QQ-plot (**Figure 6**). Although the distribution tests conducted on the simulated data predicted the distribution and shape of the data as expected, it was observed that as the number of samples decreased, the distribution of the dataset changed. This implied that histograms and qq-plots may not be the best option to use when determining the distribution of datasets with a small sample-size.

**Figure 6:**
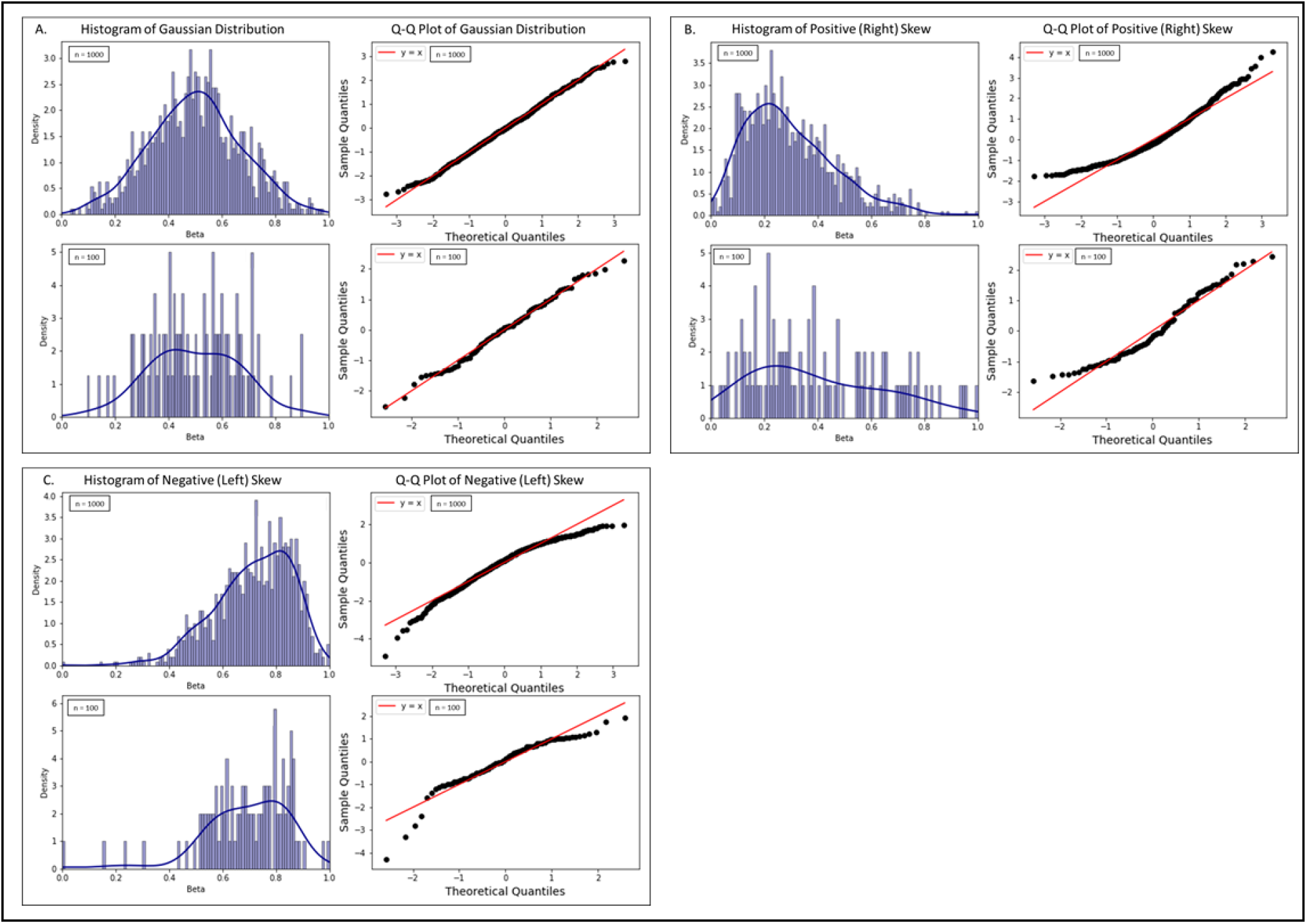
Graphical Validation of Representative Simulated Targets Showing Different Distributions using Histogram and QQ-Plot. Histograms and QQ-Plots were used to visually validate the distribution of each data set that was predicted by the custom Distribution test. **A**. Gaussian distribution shown where n = 1000 (top) and n = 100 (bottom); **B**. Positive (**Right**) Skewed Distribution shown where n = 1000 (top) and n = 100 (bottom); **C**. Negative (**Left**) Skewed Distribution shown where n = 1000 (top) and n = 100 (bottom);

### Visual validation of the “Distribution test” using histogram and QQ-Plot of TCGA data

To determine if the “Distribution test” correctly identified data sets which followed a Gaussian distribution, the methylation pattern of primary tumour and (solid) tissue normal samples for 14 TCGA cancer types across various CpG targets (covered by the HM 450K array) was screened using both a histogram and QQ-Plot (representative dataset shown in **Supplementary Figures 14 – 33**). The “Distribution test” was used to statistically predict the distribution of each dataset. By comparing both the histogram and QQ-Plot with the dScore reported by the “Distribution test”, the latter was found to be able correctly identify the overall distribution and shape of each methylation dataset.

### Influence of sample size(s) on the Overall Distribution

Non-Gaussian distributions have been frequently associated with small sample sizes due to an inadequate dispersion of the data and the frequency distribution resulting in a non-Gaussian distribution[53], [54]. To determine if the sample size has an influence on the distribution of samples at single CpG sites, the Distribution test pipeline was repeated on a random subset of tissue normal and primary tumour samples of 10 TCGA cancers (**Figure 7**). Consistent with previous literature, it was found as the number of CpG sites decreased, the overall distribution of the data would change for most CpG sites. Therefore, it is imperative to use a sufficient number of samples when performing this type of analysis. For the distribution pipeline, we recommend having a minimum of 20 samples in each group.

**Figure 7:**
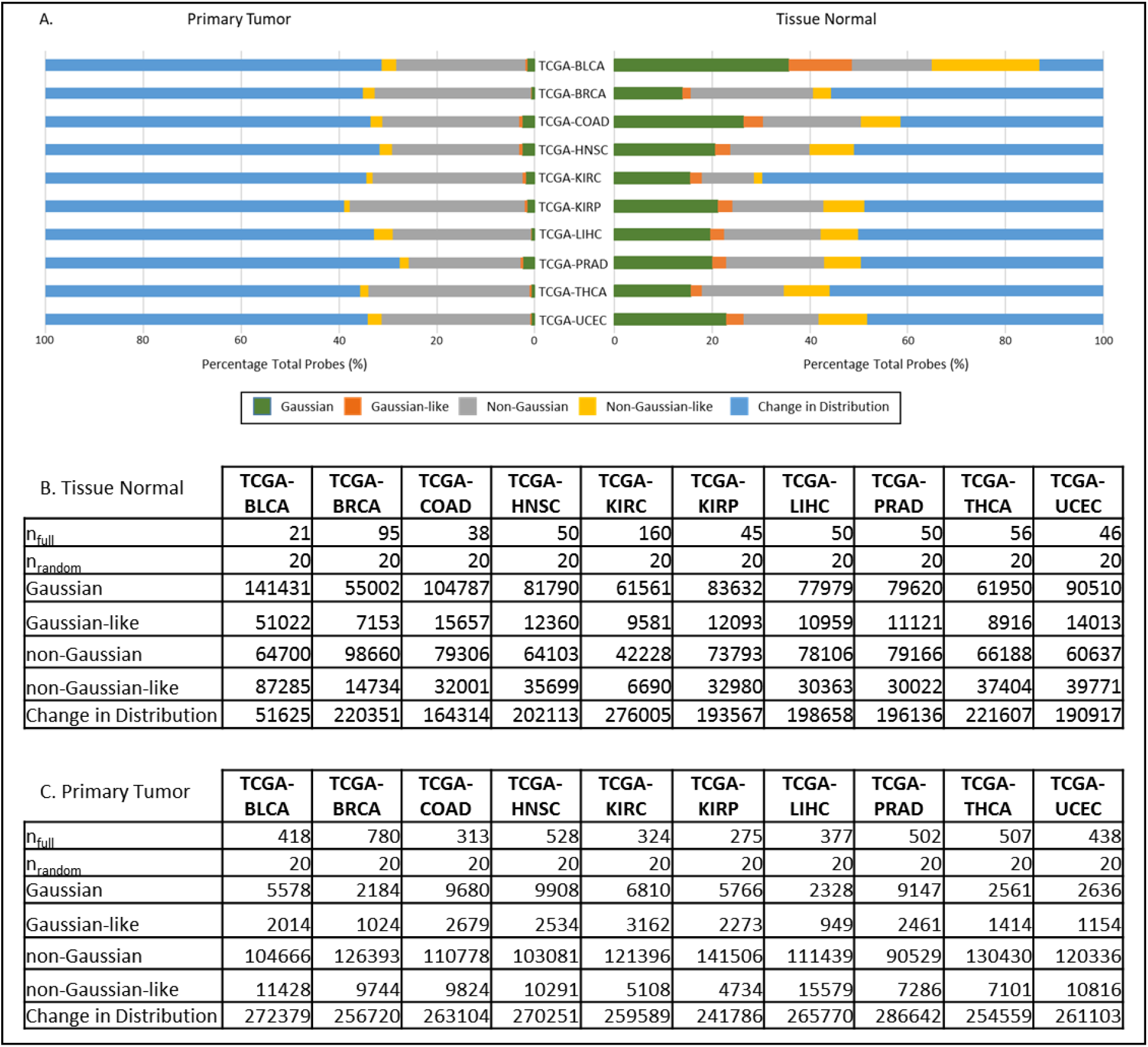
Comparison of the distribution of each CpG site when the sample size is changed. To determine if the sample size contributed to the overall distribution of each dataset, the Distribution test was repeated on a subset (n= 20) of CpG sites across 10 cancer groups where the tissue normal and primary tumor samples exceeded 20. The distribution results of the subset of data and the full CpG sample data was compared and shown in **A**. Common distributions between each subset and full dataset is shown as “Gaussian”, “Gaussian-like”, “non-Gaussian”, and “non-Gaussian-like”. In most cancer types, the percentage of probes where there was a change in distribution between the full number of samples per probe and a subset of samples appeared to be over 60 %, which suggests the sample size could play a role in the overall distributio,. Additionally, the number of matching probes for each distribution reported by the Distribution test for both Tissue Normal and Primary Tumor are shown in **B**. and **C**. respectively where nfull refers to the full number of probes across each probe, while n_random_ refers the random subset of samples from the full set.

### Comparison of Parametric and Non-parametric testing

To determine if the distribution of the methylation datasets needs to be determined prior to selecting the appropriate significance test (i.e. parametric or non-parametric) to determine the significance between two or more datasets, a parametric (t-test) and non-parametric (Mann-Whitney) test was used to determine the significance between two dataset across all probes for each project (as shown in methods). Additionally, to determine if difference in methylation pattern between datasets should be considered when using the appropriate significance test (i.e. parametric and non-parametric tests), the absolute difference in mean methylation between both primary tumour and matched (solid) tissue normal cohort data of each probe was tallied and plotted using a stacked column (**Figure 8**).

**Figure 8:**
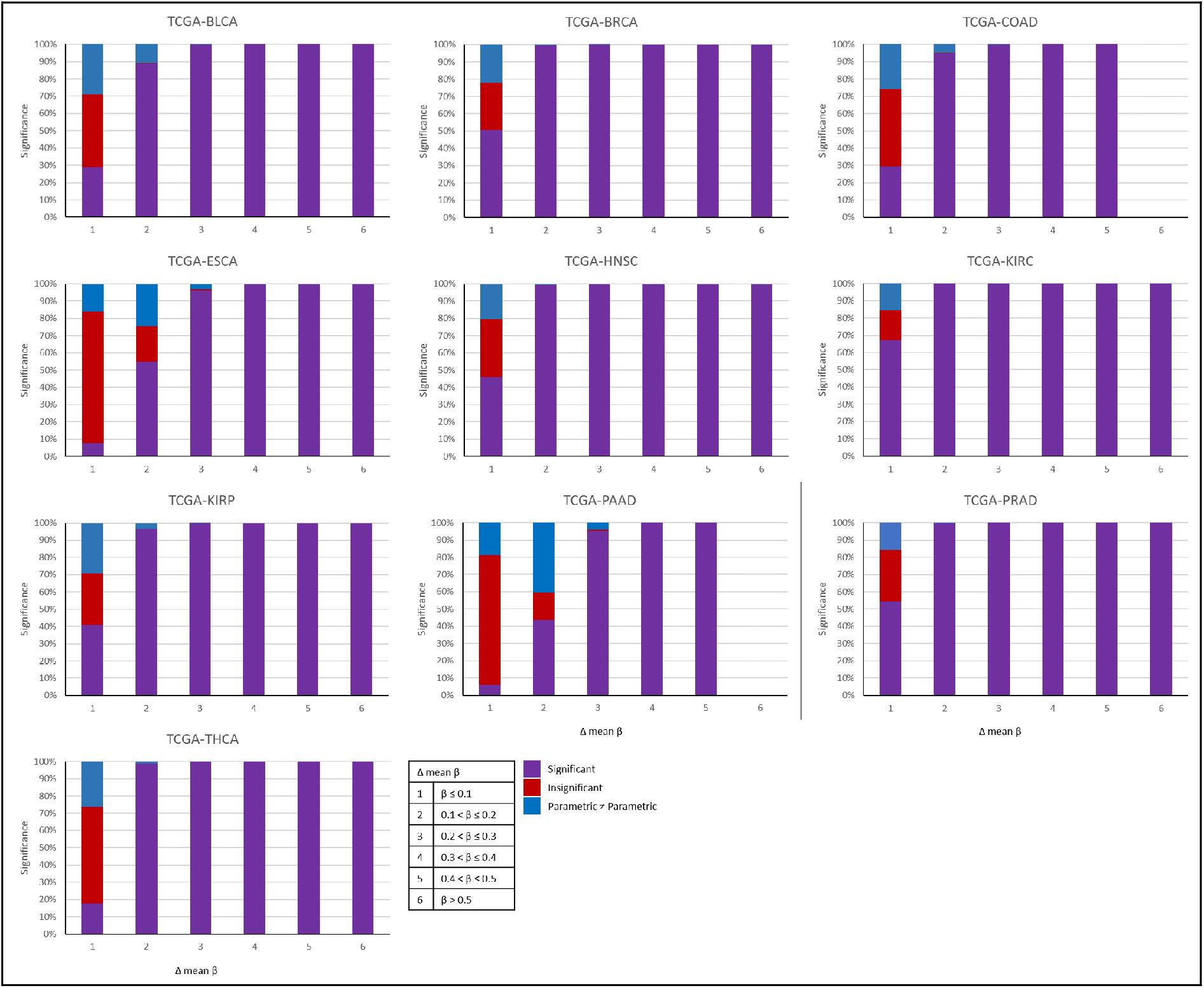
Significance testing using both parametric and non-parametric testing. Both parametric and non-parametric testing was performed on all probes across all TCGA cancer projects which had solid tissue normal samples. Ten of the fourteen cancers tested showed “significant” (purple) when tested using both the parametric and non-parametric testing regardless of the level of change in mean methylation (β). In the same ten cancers, parametric and non-parametric tests which returned “insignificant” (red) were only present in CpG sites with a mean difference of less than 10 – 20 % methylation.

For this analysis, only datasets which followed a Gaussian (dScore = 5), or non-Gaussian distribution (dScore = 0) were considered. From this analysis, it was found that both the parametric t-test and the non-parametric Mann-Whitney test used showed significance between solid tissue normal and primary tumour across the majority of cancer types when the absolute difference in mean β is less than 0.1 – 0.2 (i.e. 10 – 20 % methylation). This was consistent with findings from past literature where biomarkers with a difference in mean methylation of 10 – 20 % was considered as “significant” biomarkers with potential for future testing [18], [19], [22], [30].

The four cancers which showed significant and non-significant probes when tested using parametric and non-parametric testing across all changes in mean methylation was shown in **Supplementary Figure 34,** and are cancers that are associated with the lung, liver and uterine. Furthermore, to determine if the distribution of the two data sets influenced the outcome of the significance testing, the change in distribution between each set of data was also observed. It was found that for most of the cancers tested, as the level of mean difference in beta increased, the significance between the sample cohorts increased.

### Biomarker parsing using feature selection – Quartile Parsing

“Quartile Parsing” is used an alternative method to identify biomarkers which are differentially-methylated between different cancers. In this study we used the difference in quartiles between two different cancers (TCGA-BRCA and TCGA-PRAD). For a biomarker to be “hypermethylated”, the 1^st^ quartile (Q1, 25^th^ percentile) of the primary tumour needs to be above 0.7 (70%), while the 3^rd^ quartile (Q3, 75^th^ percentile) needs be below 0.3 (30%). In contrast, for hypomethylated biomarkers, the 3^rd^ quartile (Q3, 75^th^ percentile) needs to be below 0.3 (30%) and the 1^st^ quartile (25^th^ percentile) needs to be above 0.7 (70%). This method was compared to the significance testing method (**Figure 9**), where biomarkers that has a p value of below 0.01 were considered to be potential biomarkers. Initial analysis using clustering and heatmap suggests the “Quartile Parsing” method is adequate as an initial analysis to find differentially-methylated biomarkers that have distinct methylation patterns.

**Figure 9:**
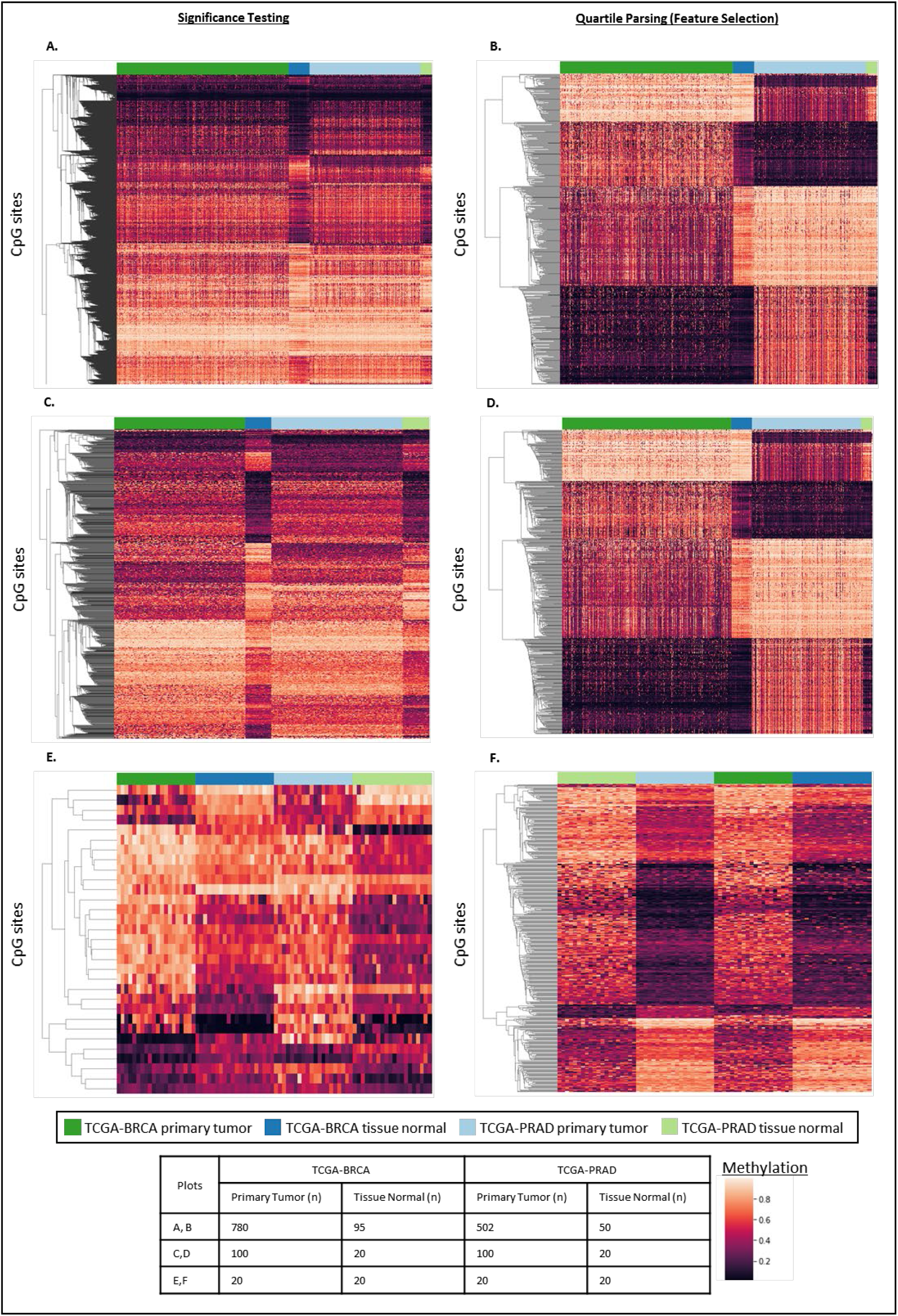
Mining for differentially-methylated biomarkers between TCGA-BRCA and TCGA-PRAD using significance testing and quartile parsing. Heatmaps generated from biomarkers identified using significance testing, where **A**) the full set of samples was used, (**C**) 100 random primary tumour samples and 20 random tissue normal samples from each cancer was used, (**E**) 20 random primary tumour samples and 20 random tissue normal samples from each cancer was used. Heatmaps generated from biomarkers identified using “Quartile Parsing”, where the (**B**) the full set of samples was used, (**D**) 100 random primary tumour samples and 20 random tissue normal samples from each cancer was used, (**F**) 20 random primary tumour samples and 20 random tissue normal samples from each cancer was used. From the heatmaps generated, the quartile parsing appeared to be able to parse biomarkers that present a clear difference in methylation between the two cancers tested.

## Discussion

In recent years, there has been an explosion of data to support the importance of DNA methylation in cancer. The development of the Infinium HumanMethylation450 Bead-Chip® (HM450) array has allowed researchers to perform high-throughput methylation profiling on more than 450, 000 CpG targets throughout the genome [55]. The ability of the platform to quickly, and affordably perform genome-scale methylation analysis aligns with the needs for large-scale projects such as The Cancer Genome Atlas (TCGA) to examine and curate the methylation profile of hundreds of samples simultaneously. The TCGA project have released multi-omic data on over 20,000 primary cancer and matched normal samples spanning 33 different cancer types, with the goal to improve our ability to diagnose, treat, and prevent cancer [56], [57]. The methylation data is publicly-available as β files, where the level of fluorescence for methylated (M,0) and unmethylated (U, 0) probes at each site have already been filtered and normalised for downstream analysis [58]. Currently, a number of software packages and methods have been developed to process and streamline the analysis of HM450 data (such as *IMA, Lumi, Minfi* etc) [39], [59]. Generally, for large datasets such as these generated by the HM450 arrays, most programs will use non-parametric statistical tests (e.g. Wilcoxon’s signed rank test or Mann-Whitney test etc) to determine the statistical significance between samples and between groups (i.e. statistical significance is observed if the p value(s) is below a certain threshold e.g. < 0.05), as non-parametric tests generally does not make assumptions about the shape or distribution of the data. In response, we established a “Distribution test” pipeline which determines the distribution and shape of a given methylation data set (in beta values), and from this selects the optimal inferential statistics to use on each dataset. The purpose of this pipeline is to enable researchers to make better choices when selecting the most appropriate statistical methods on each dataset.

Histograms and QQ-Plots are the most convenient methods to determine the distribution or shape of a dataset, and was used in this article to determine if the Distribution test pipeline was correctly calling the distribution and shape of each probe-wise methylation dataset. However, due to the large number of datasets handled through methylation studies, it is not logical to use visual validations to determine the distribution of the data. Therefore, in this study, a series of distribution analysis algorithms was incorporated into the custom “Distribution test” in order to determine the distribution, shape and skewness of the methylation datasets and report it as a “distribution Score” (dScore). In order to determine the robustness of this pipeline in determining the distribution and shape of each cohort of methylation data, different sets of simulated β values with different distributions were visualised using both a histogram and QQ-plot (representative data in **Figure 7**, and other representative data in **Additional Figures 1 - 20**). The results suggested that the combination of tests used in the “Distribution test” was effective in describing both the distribution and shape of the methylation datasets. However, visual validation of data distribution becomes less accurate as the sample size decreases, leading to a false interpretation of the overall distribution.

The Distribution test pipeline was used to determine the distribution of each cohort of samples (primary tumour and matched (solid) tissue normal) across individual probes (i.e. CpG sites) of the HM450K array of 14 cancers from the TCGA project. This analysis returns a distribution score (dScore) for each group of samples, that indicates whether the data followed a Gaussian or non-Gaussian distribution. Furthermore, we determined if there any implications of using parametric and non-parametric tests on datasets where there is a shift in distribution between each cohort of specific CpG sites. This enabled the selection of the optimal inferential test to screen for CpG sites that showed statistical significance between the samples, which when visualised using a semi-supervised clustering method, differential methylation can be observed between the samples used in this analysis (**Figure 9 A, C, E**). Finally, we proposed “Quartile Parsing” as an alternative method to screen for differentially-methylated biomarkers in future projects. Based on feature selection, this pipeline screened for CpG sites that showed distinct differences in methylation by comparing the differences in methylation at different quartiles. Visualisation of biomarkers filtered using this method, highlighted biomarkers that showed distinct changes in methylation between different tissues or tumours (**Figure 9 B, D, F**). Therefore, this pipeline was introduced as an alternative method to screen for biomarkers that presented a clear contrast in methylation between different tissues or tumour samples.

## Conclusions

In conclusion, the Illumina Human Methylation 450 Bead Chip array is a high-throughput platform that is popularly used to screen for diagnostic and prognostic biomarkers for cancer. A number of softwares have been released for processing and analysing HM450K array data. These softwares often use different statistical methods to determine the “significance” between and within sample populations. A survey of past literature has found that statistical significance is often used as the only analysis in screening for differentially methylated biomarkers. However, this is not always appropriate as these methods often highlighted biomarkers with minimal differences in methylation (i.e. <10 – 20%). Choosing the right statistical methods depends largely on the distribution of the dataset (i.e. Gaussian or non-Gaussian), an avenue, which is rarely explored in the epigenetic space. In this study, we explored the implications of using different statistical methods (i.e. parametric and non-parametric) on methylation datasets with varying distributions.

By exploring the distribution of each group of samples across individual CpG sites with our custom “Distribution test”, this study showed that both parametric and non-parametric tests showed biomarkers with 10 – 20 % absolute difference in mean methylation between the primary tumour and its matching (solid) tissue normal samples from ten of the fourteen TCGA projects used. The Distribution test pipeline was introduced to firstly predict the distribution and shape of input datasets, and select the optimal inferential statistical method for predicting the differences in methylation between sample groups. Additionally, we introduce the use of “Quartile Parsing” as an alternative method to screen for biomarkers with distinct methylation patterns between tissue normal and cancers of interest.

