## Supplementary figures and images for "An analytical pipeline for DNA Methylation Array Biomarker Studies"

### Supplementary Figure 1

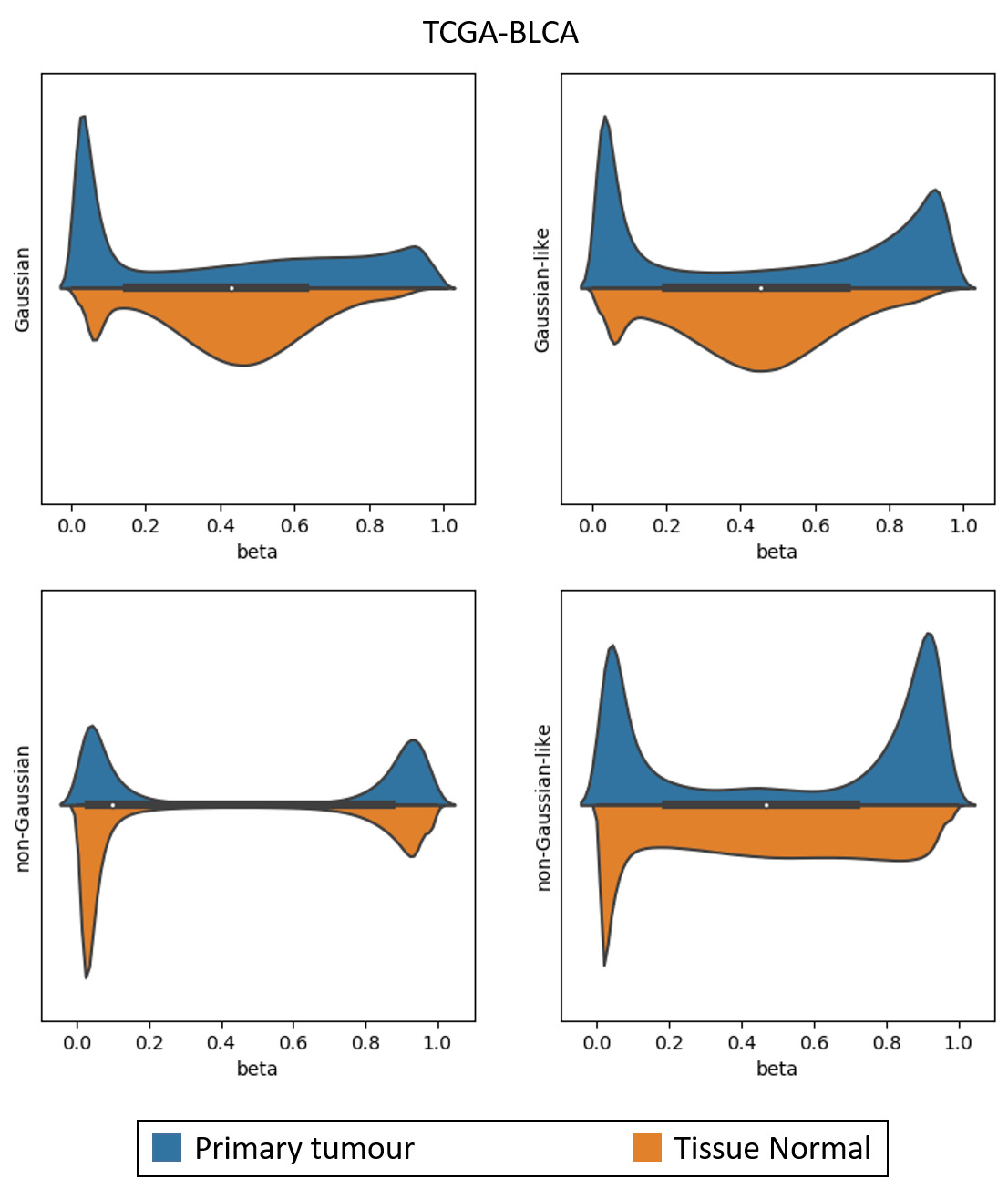

### Supplementary Figure 2

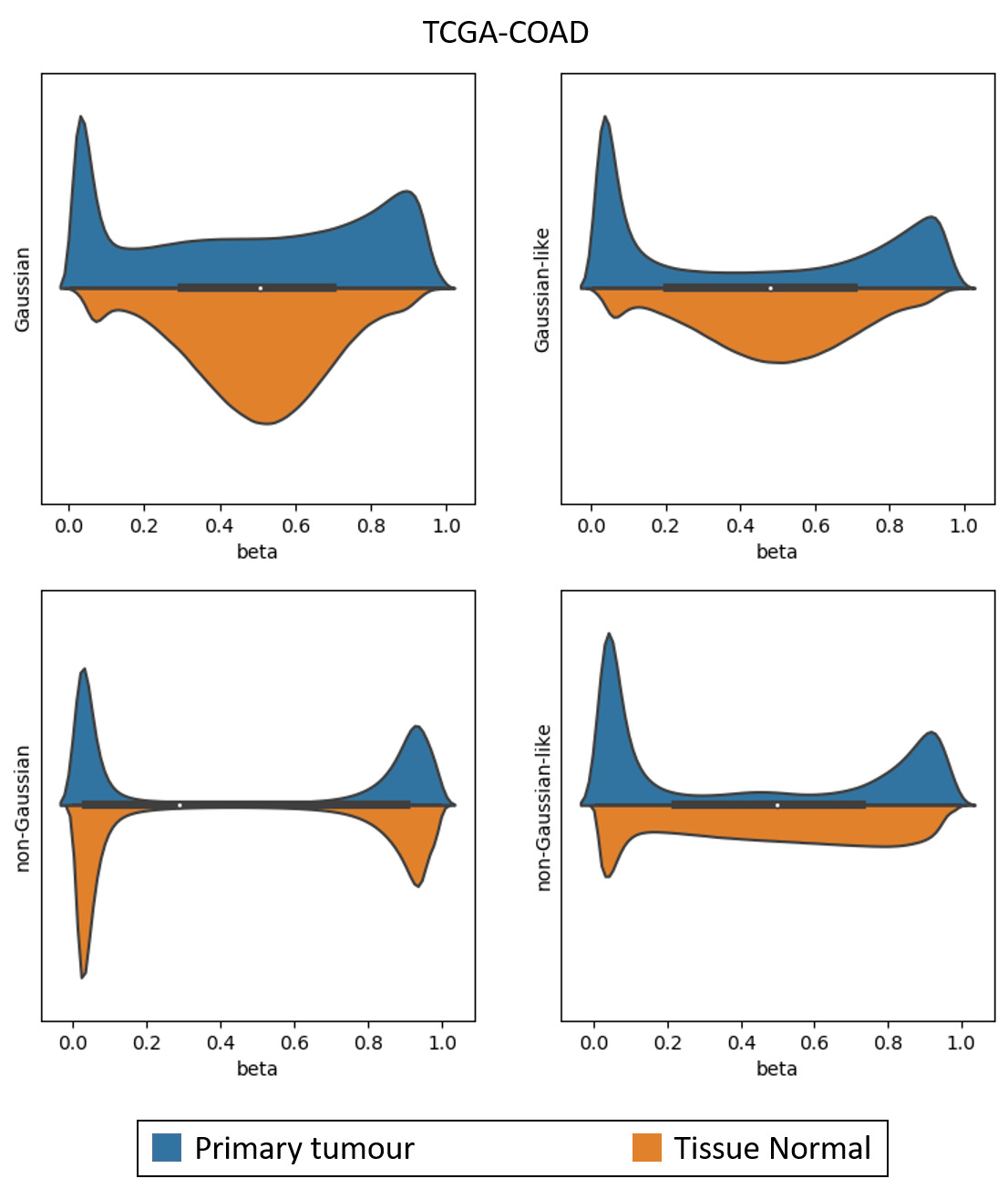

### Supplementary Figure 3

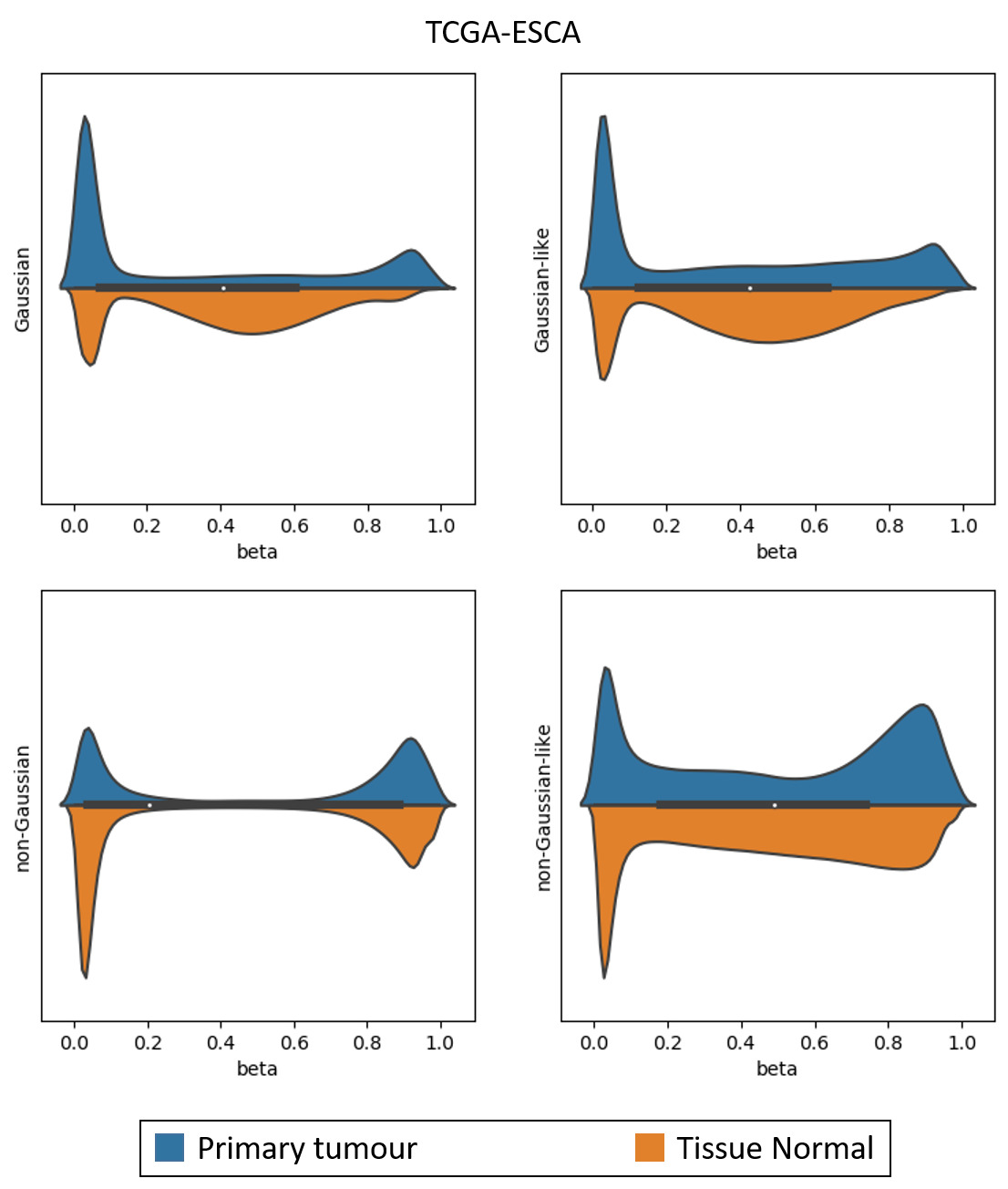

### Supplementary Figure 4

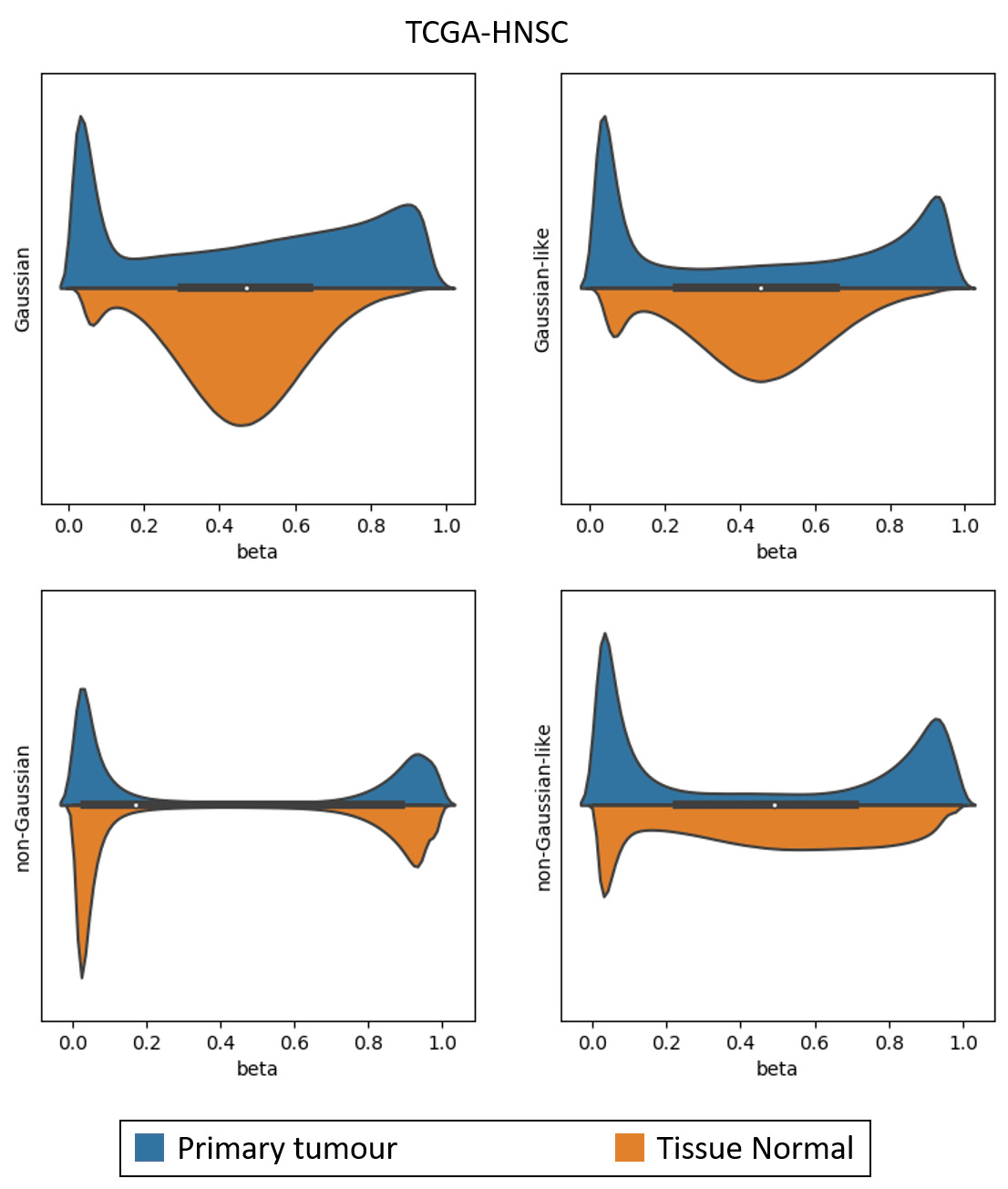

### Supplementary Figure 5

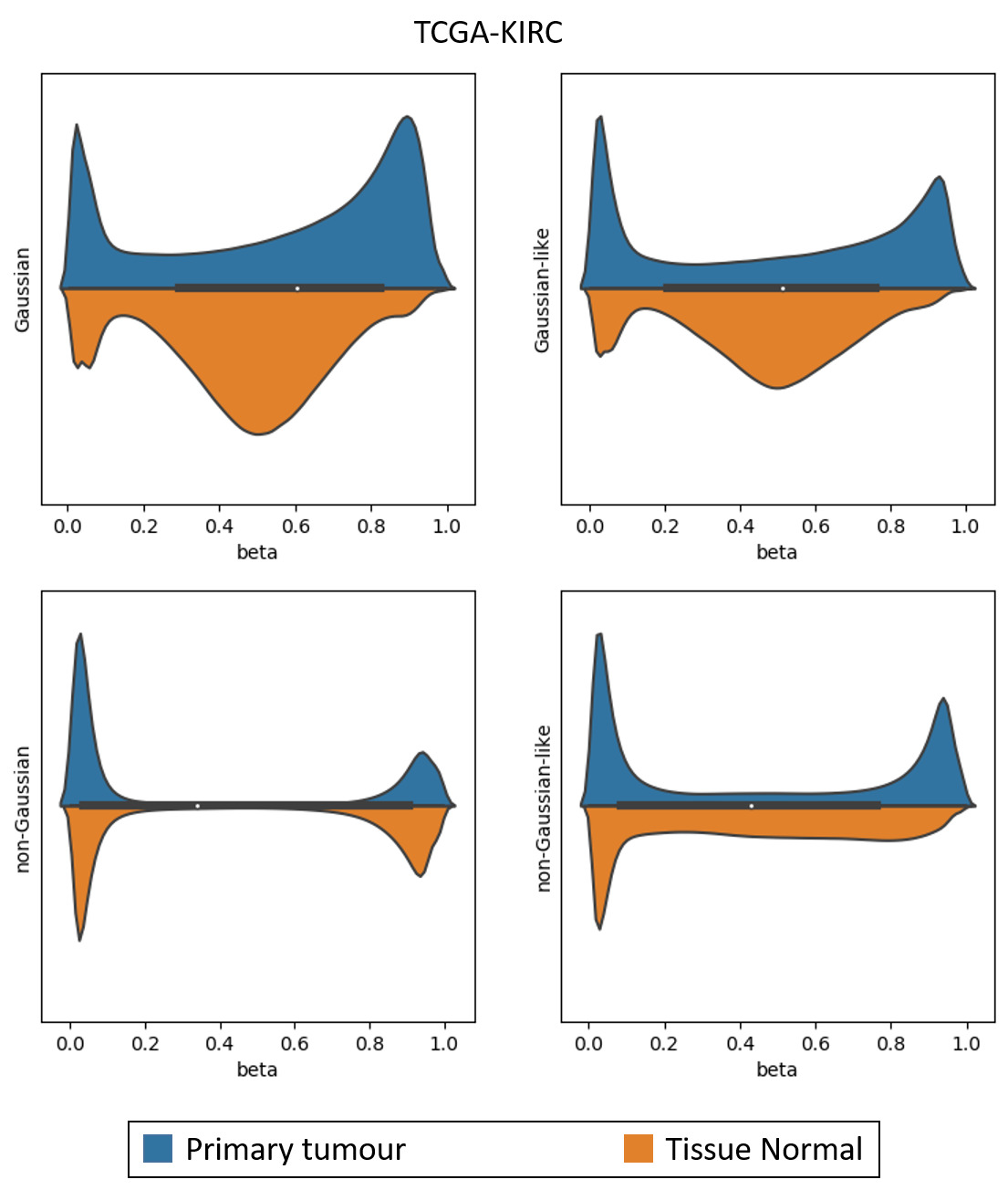

### Supplementary Figure 6

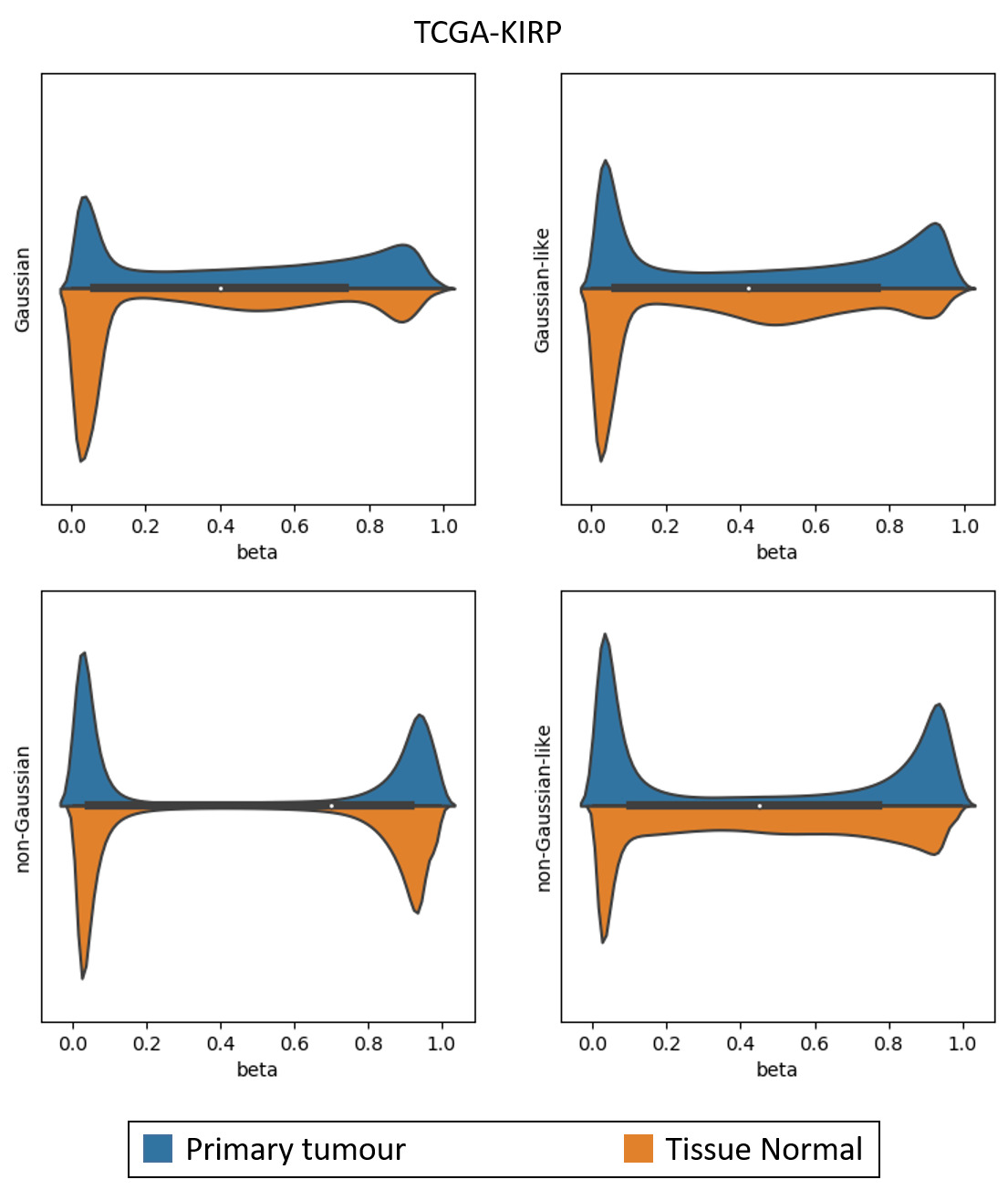

### Supplementary Figure 7

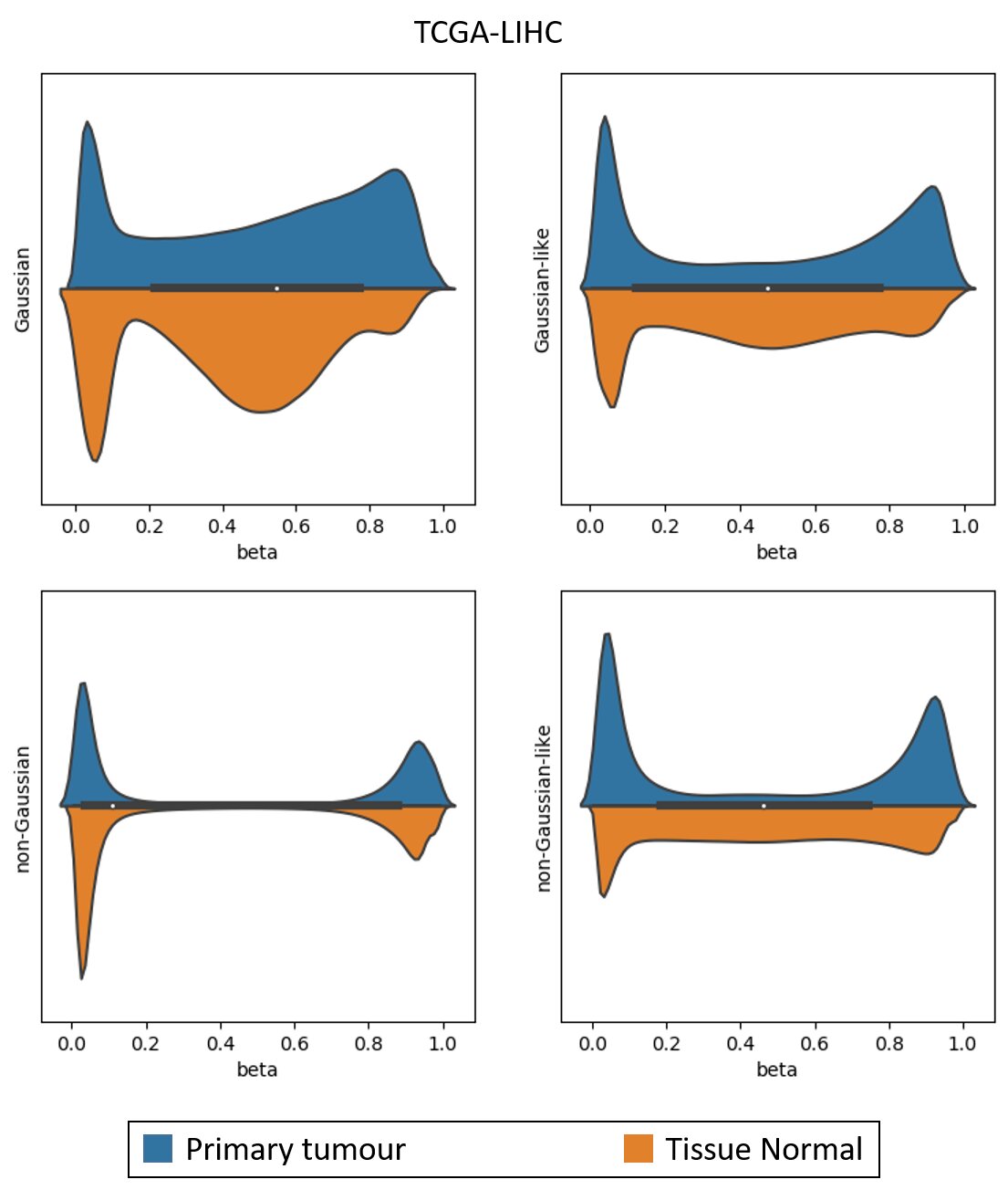

### Supplementary Figure 8

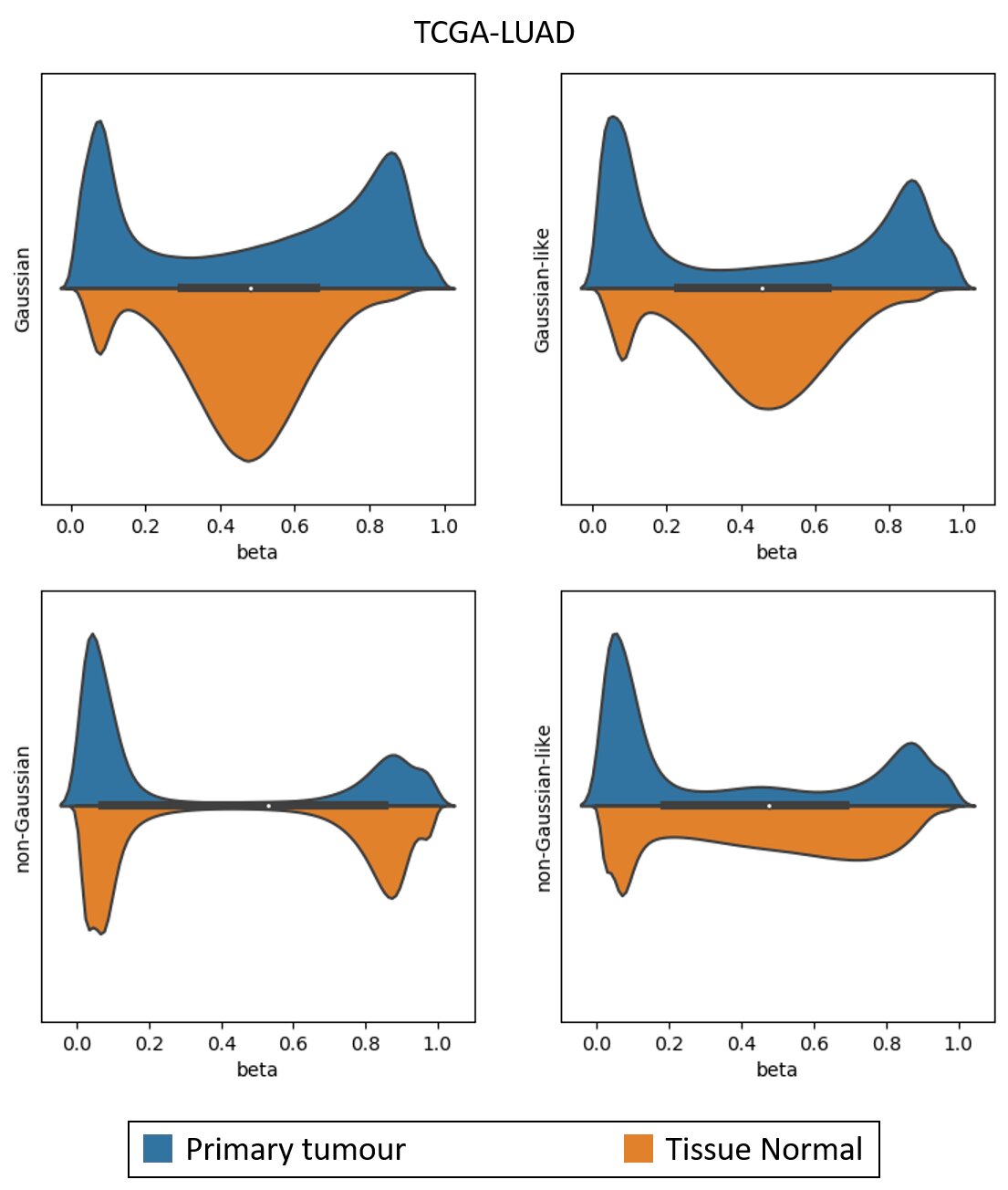

### Supplementary Figure 9

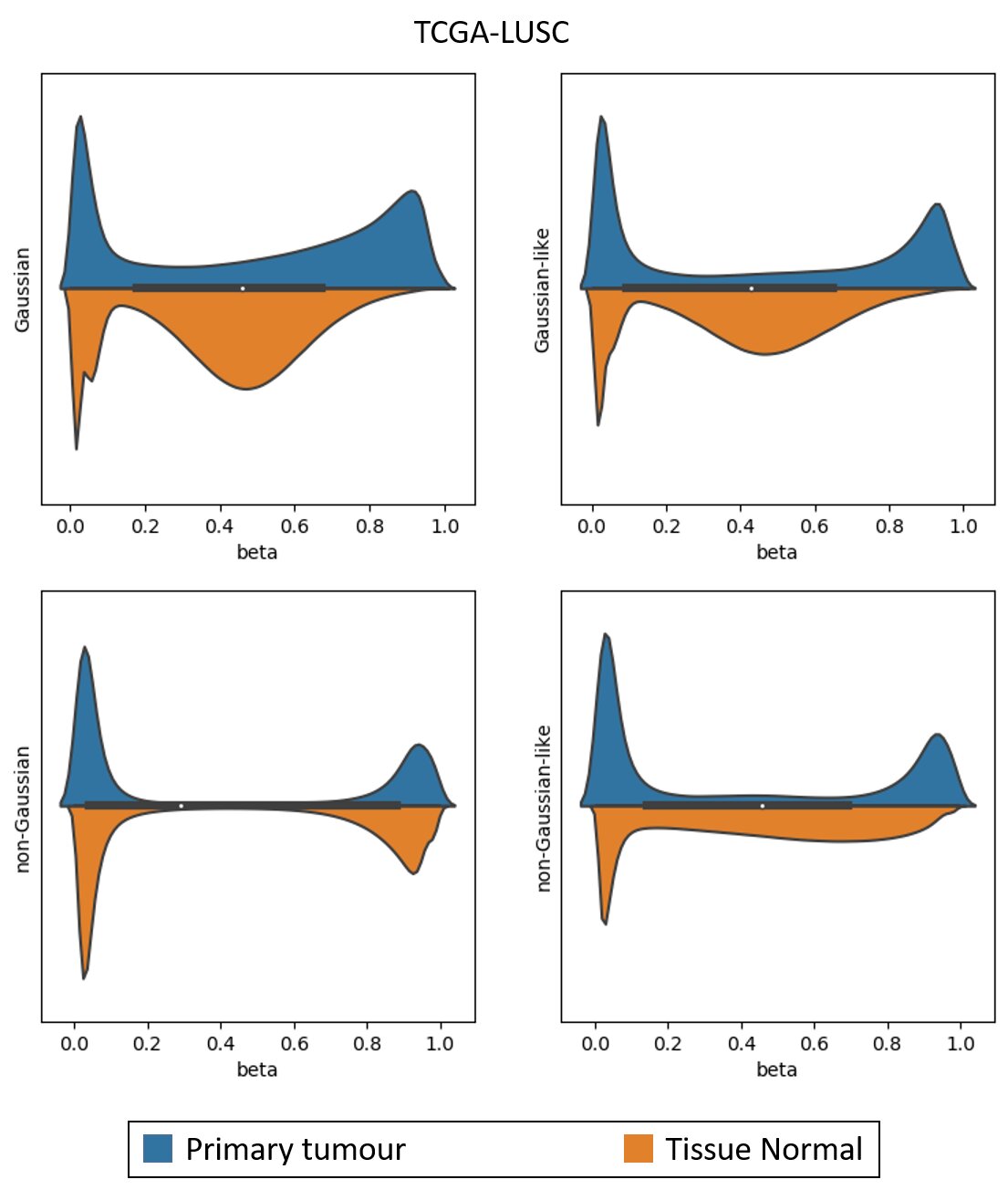

### Supplementary Figure 10

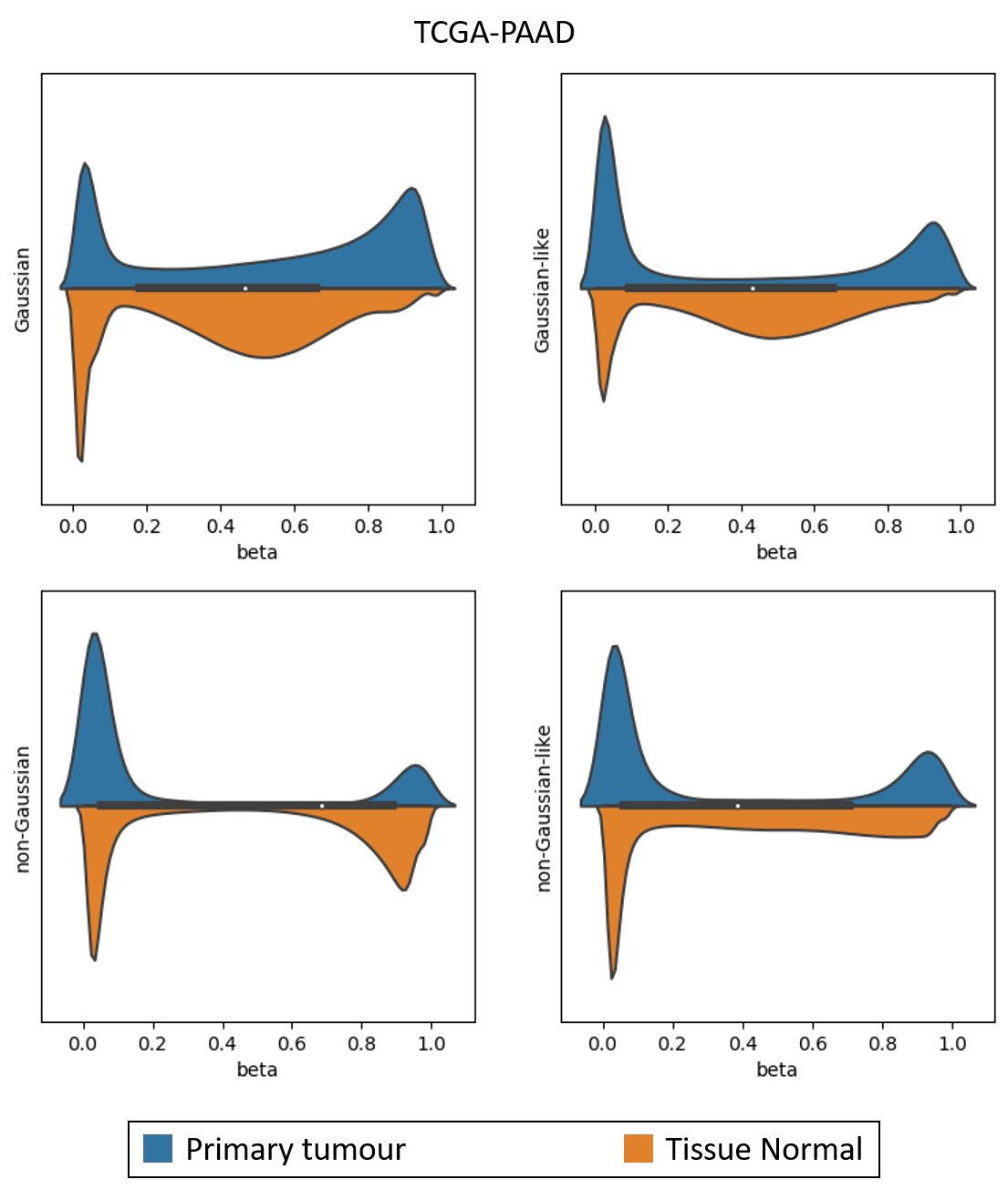

### Supplementary Figure 11

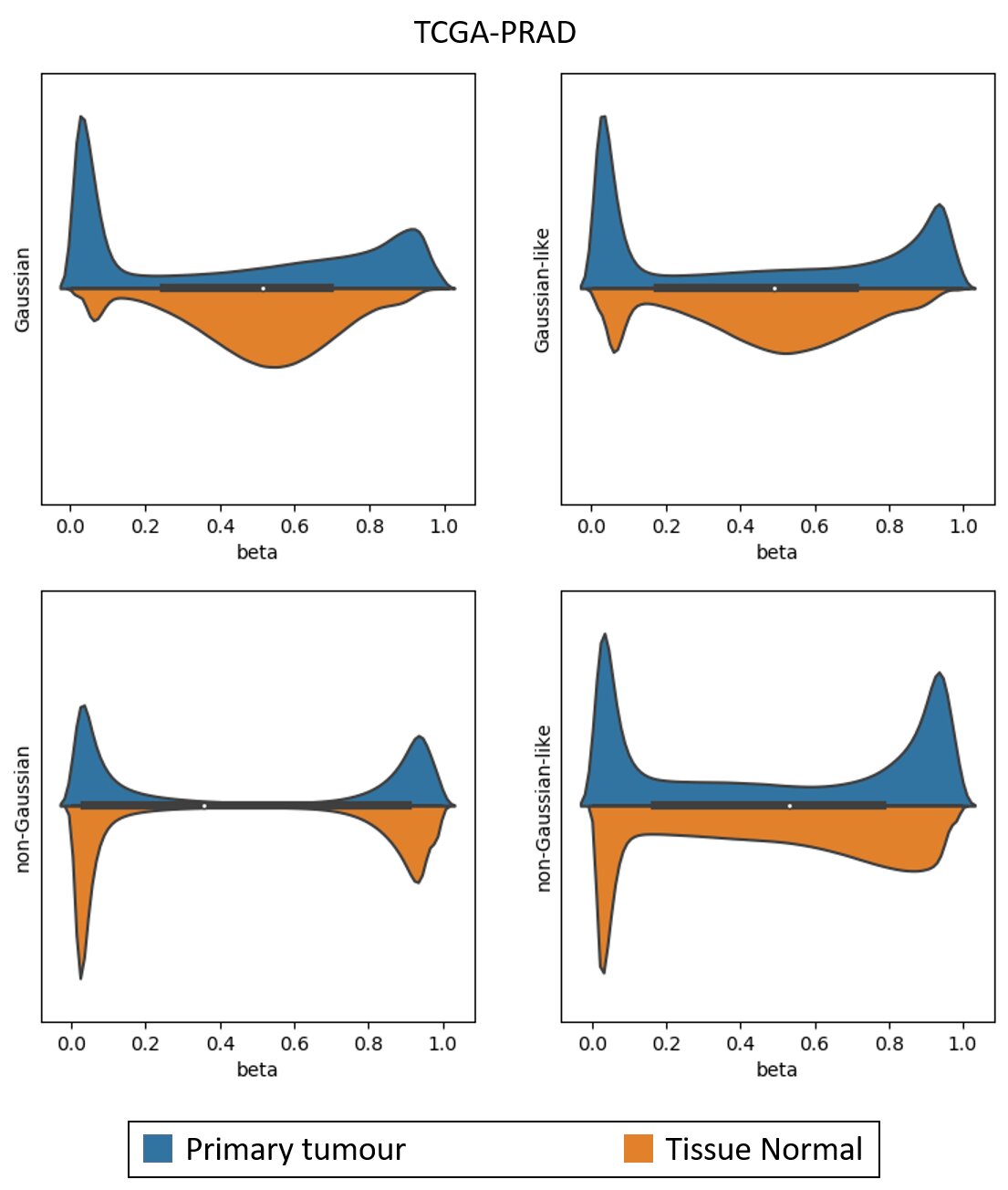

### Supplementary Figure 12

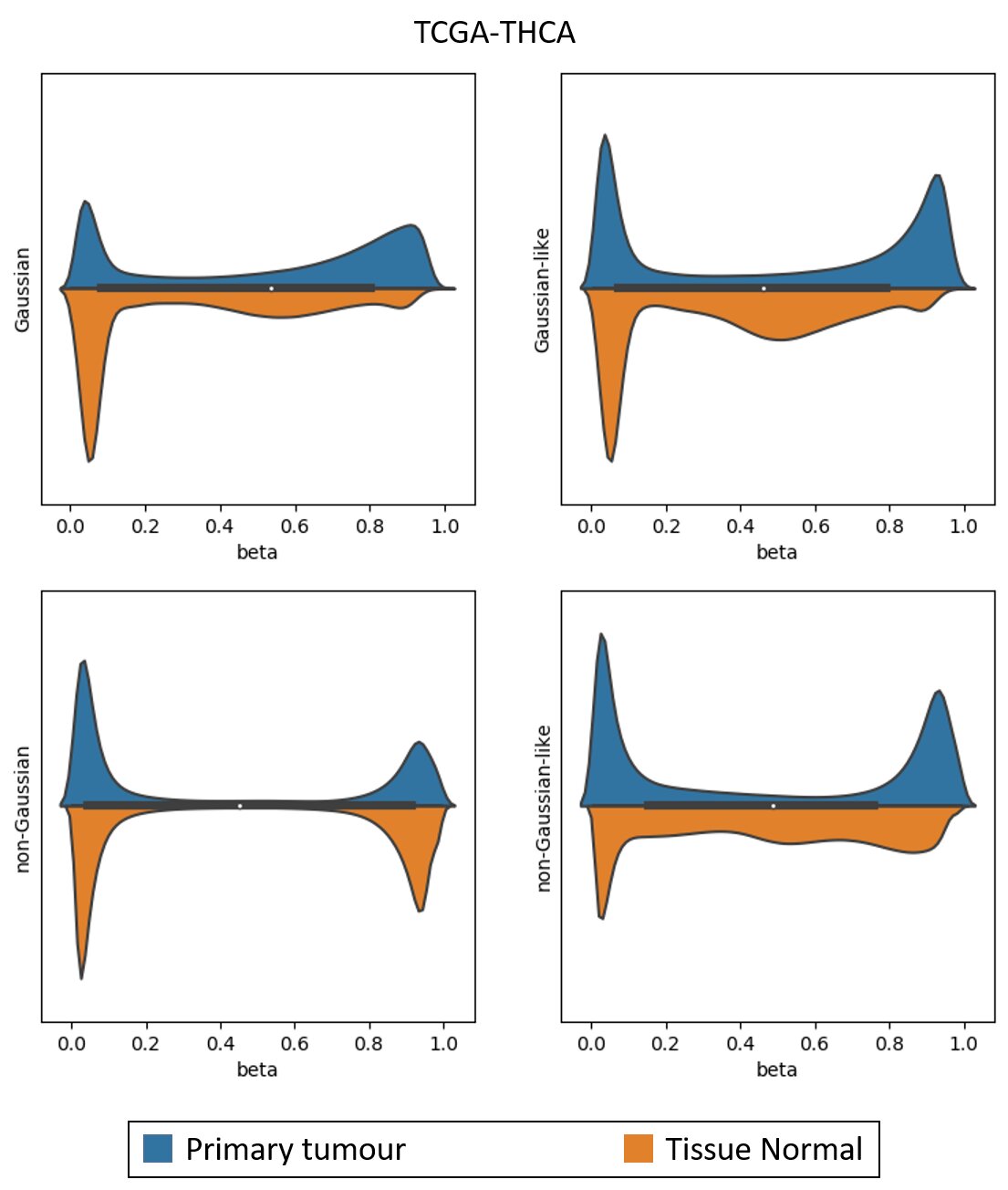

### Supplementary Figure 13

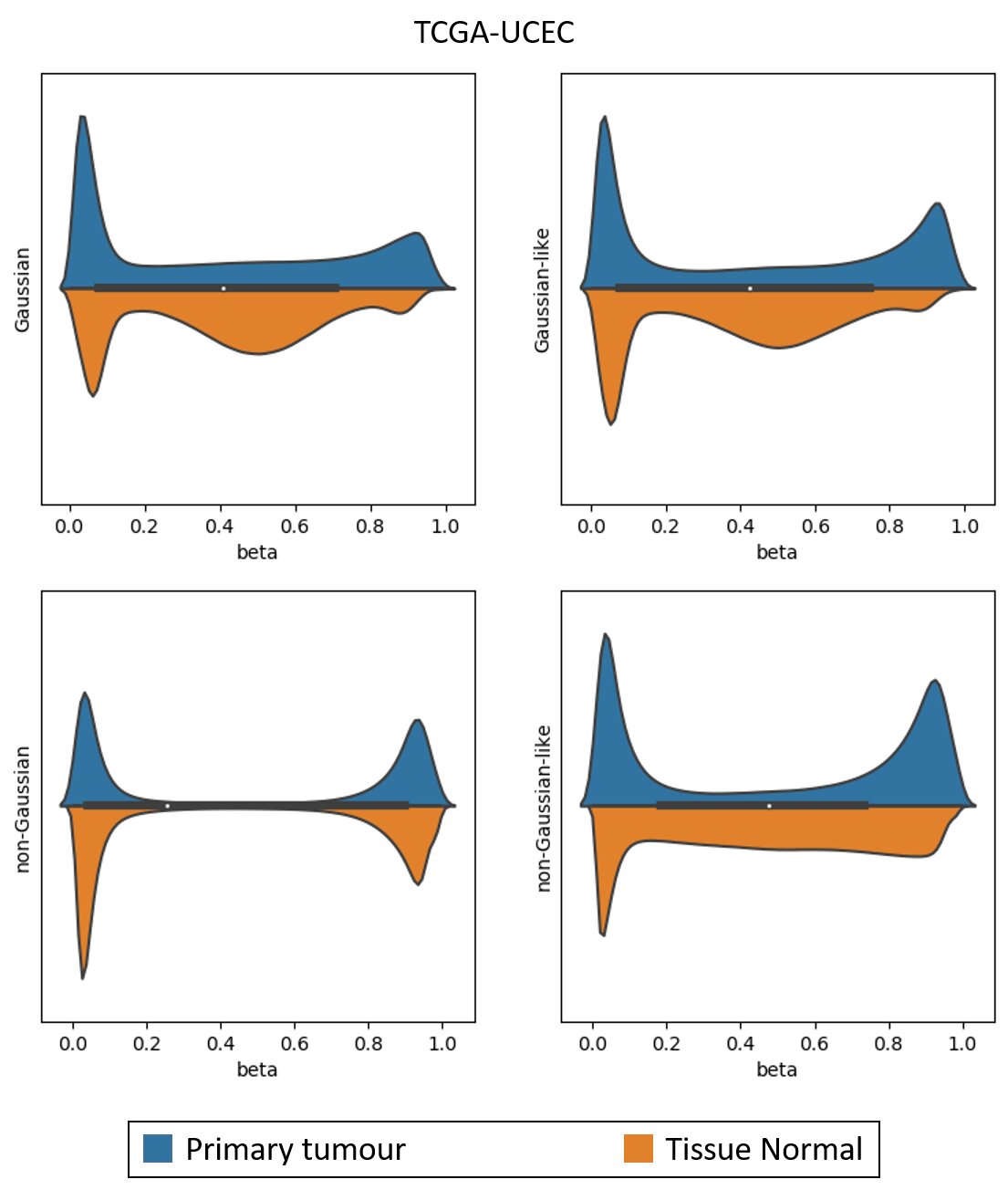

### Supplementary Figure 14

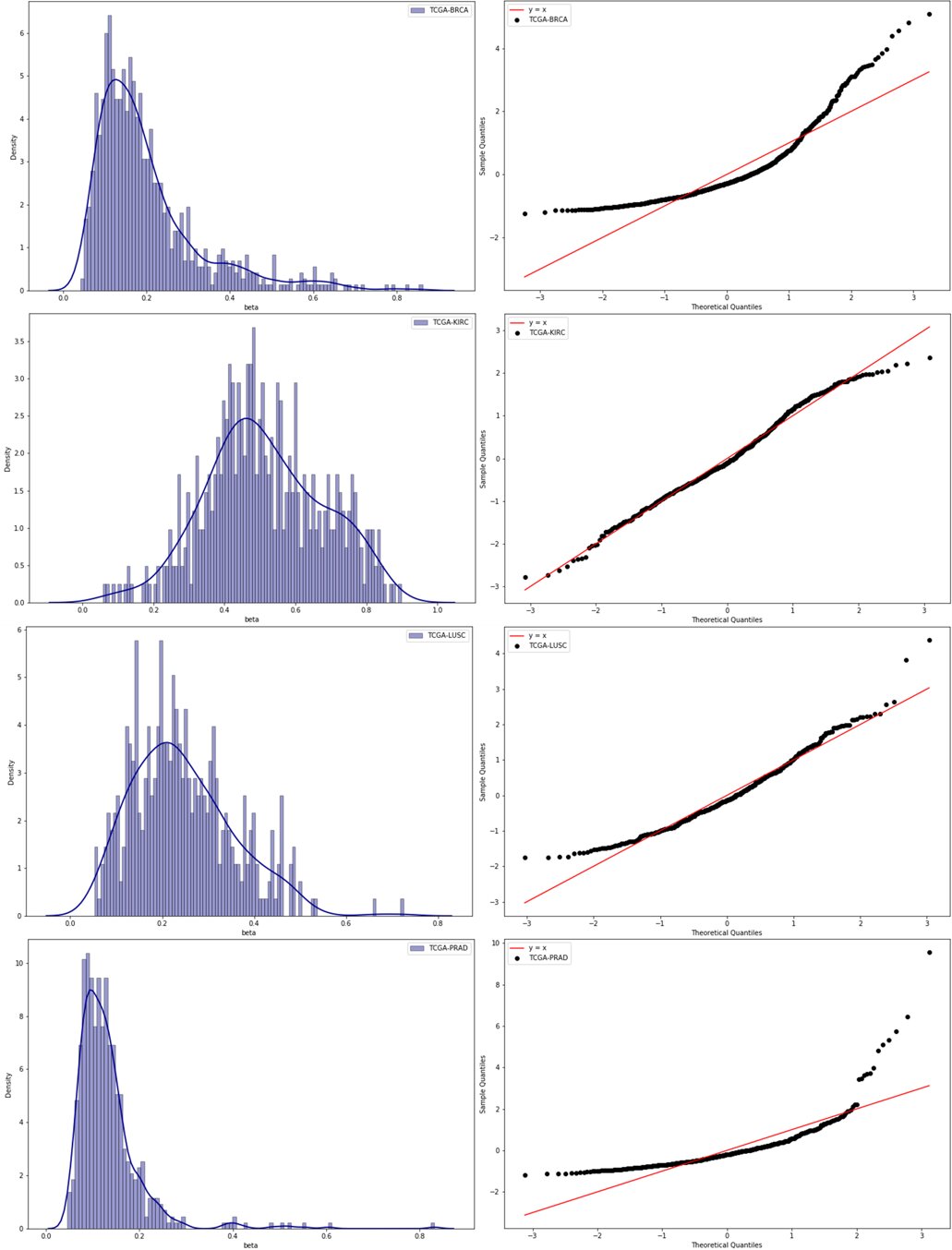

### Supplementary Figure 15

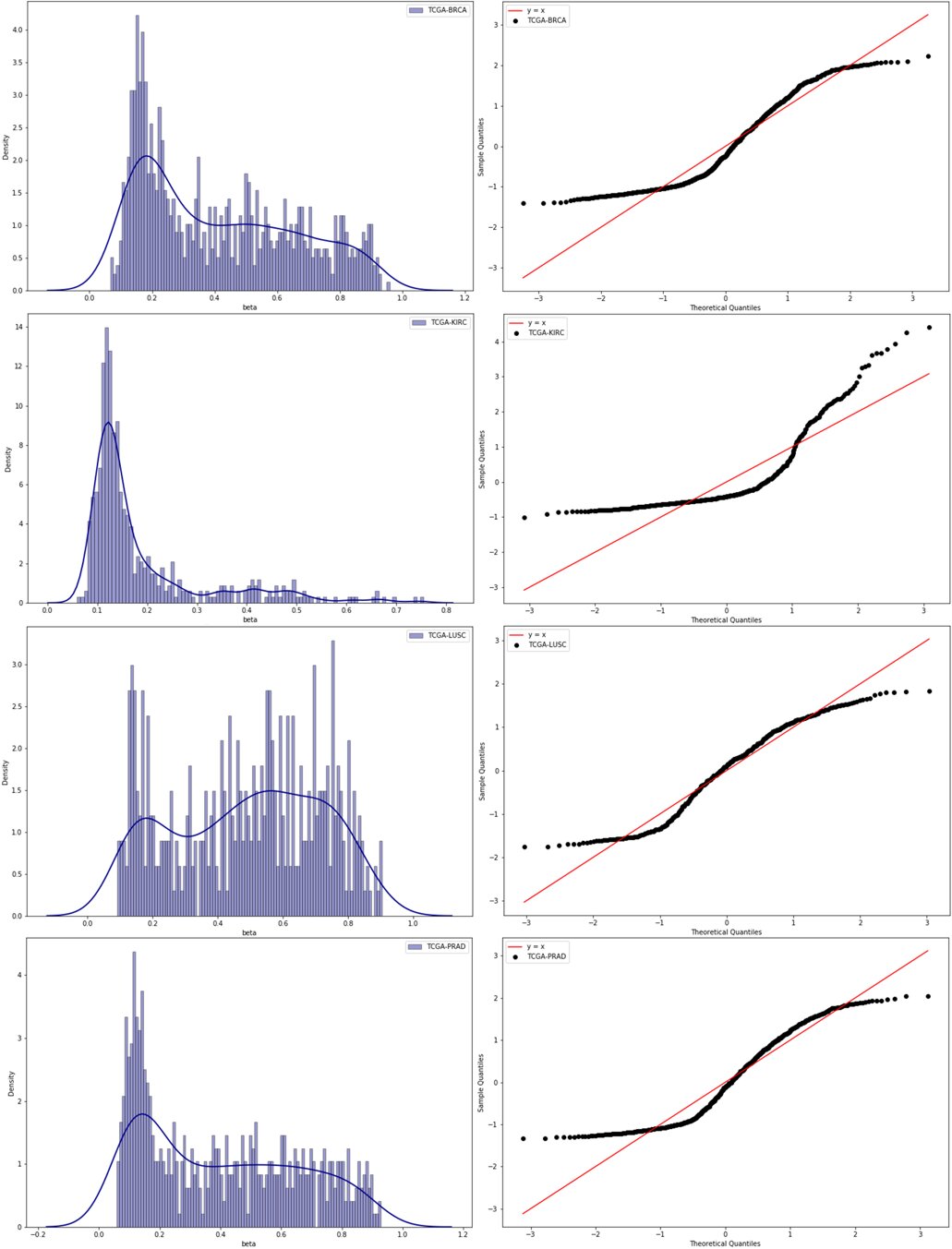

### Supplementary Figure 16

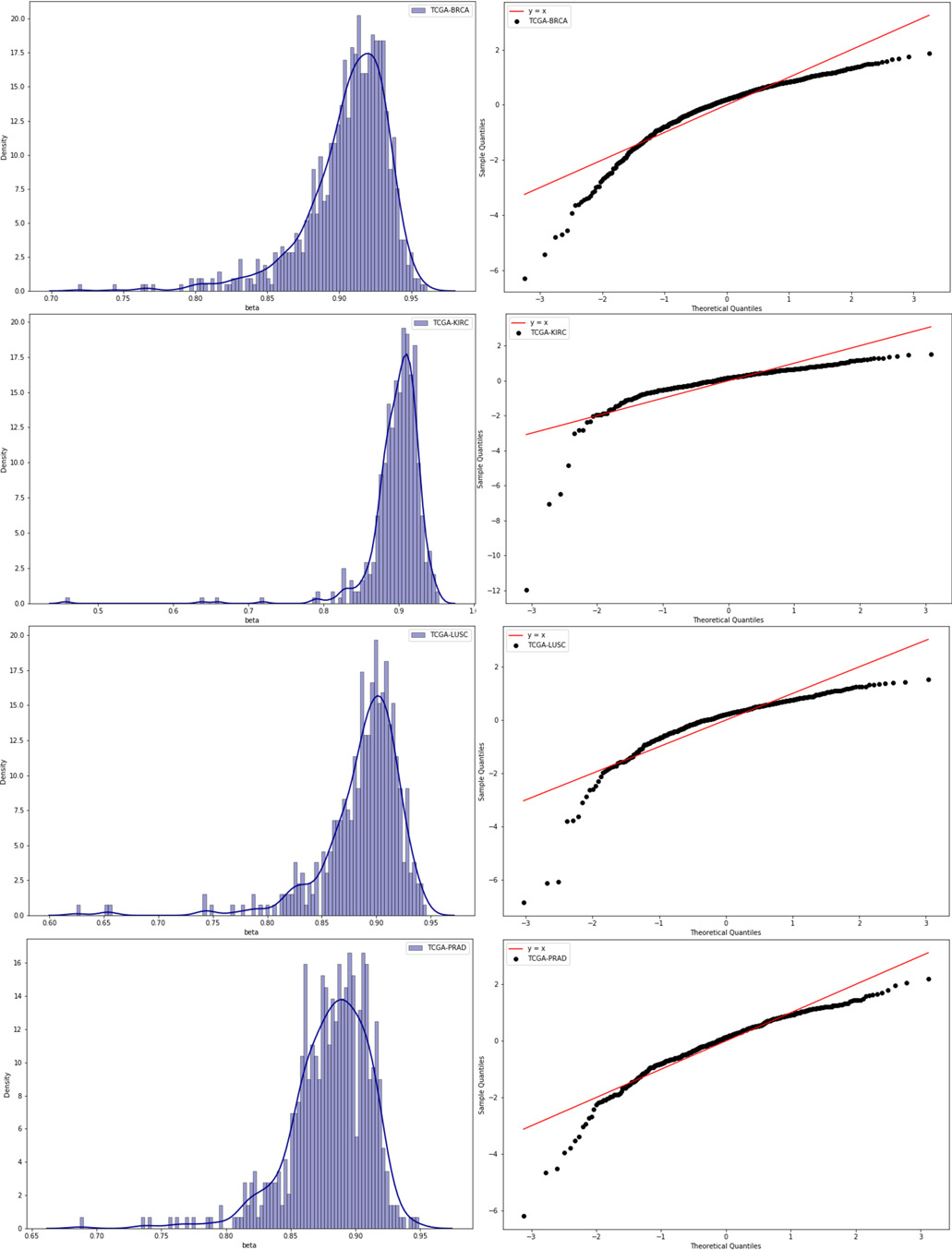

### Supplementary Figure 17

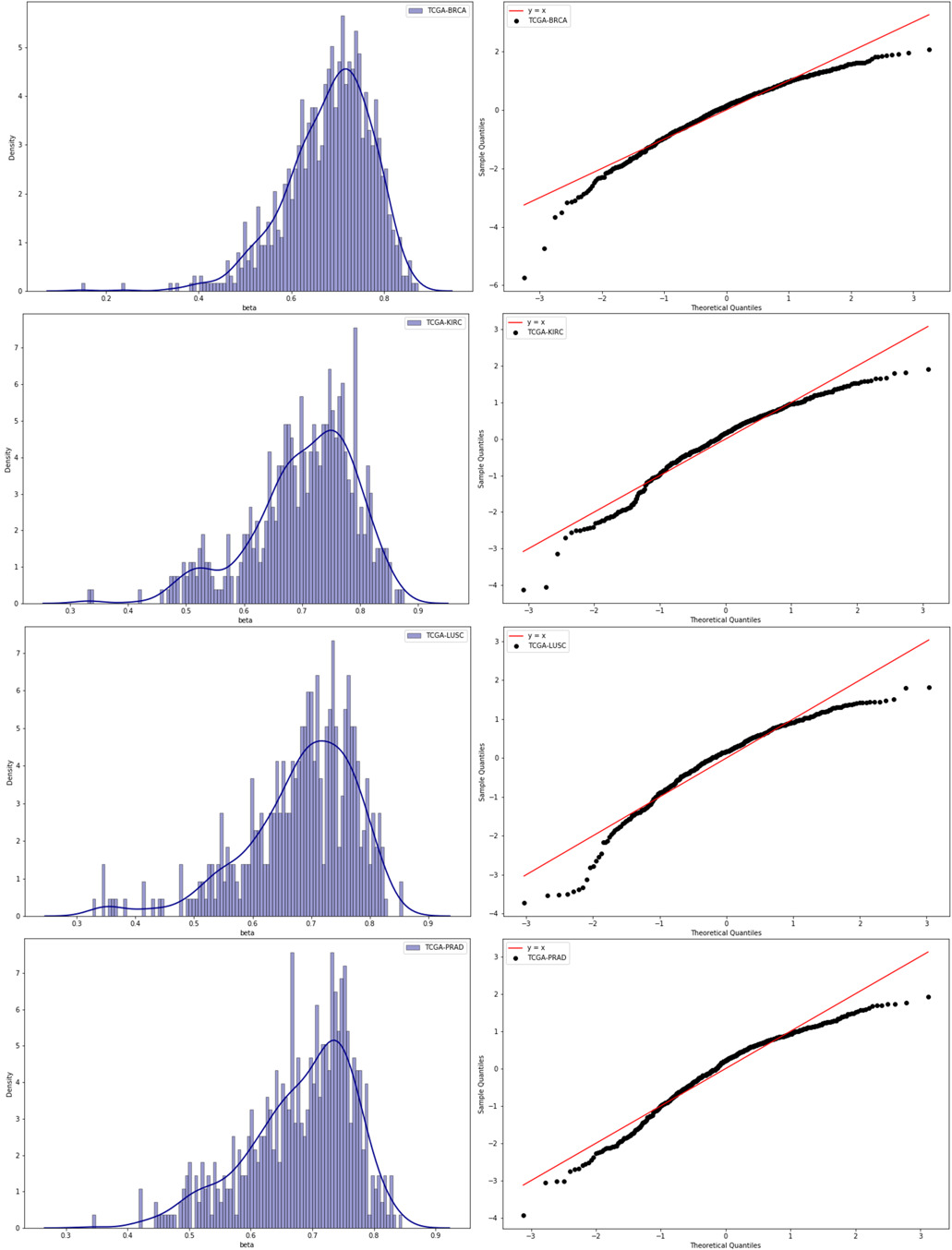

### Supplementary Figure 18

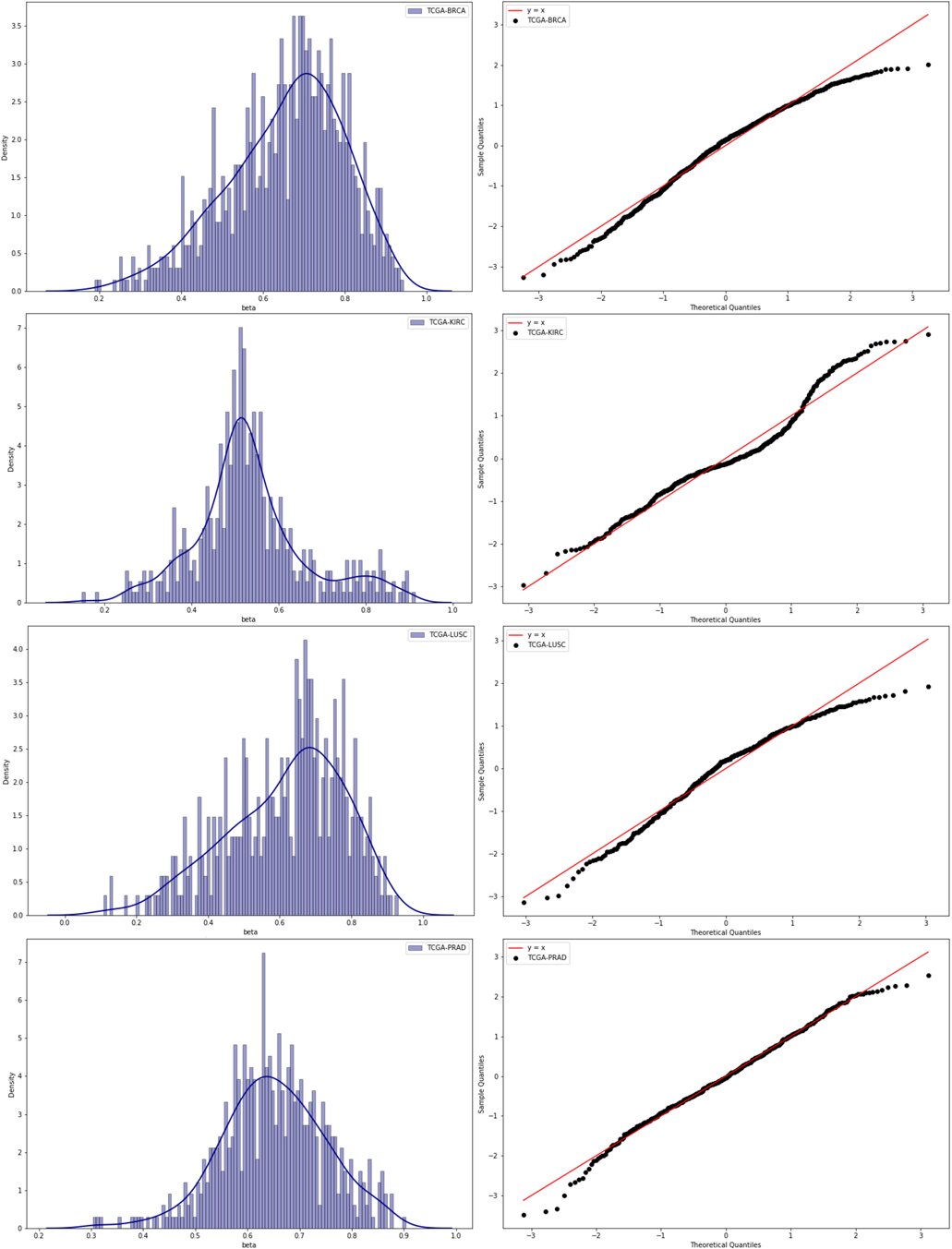

### Supplementary Figure 19

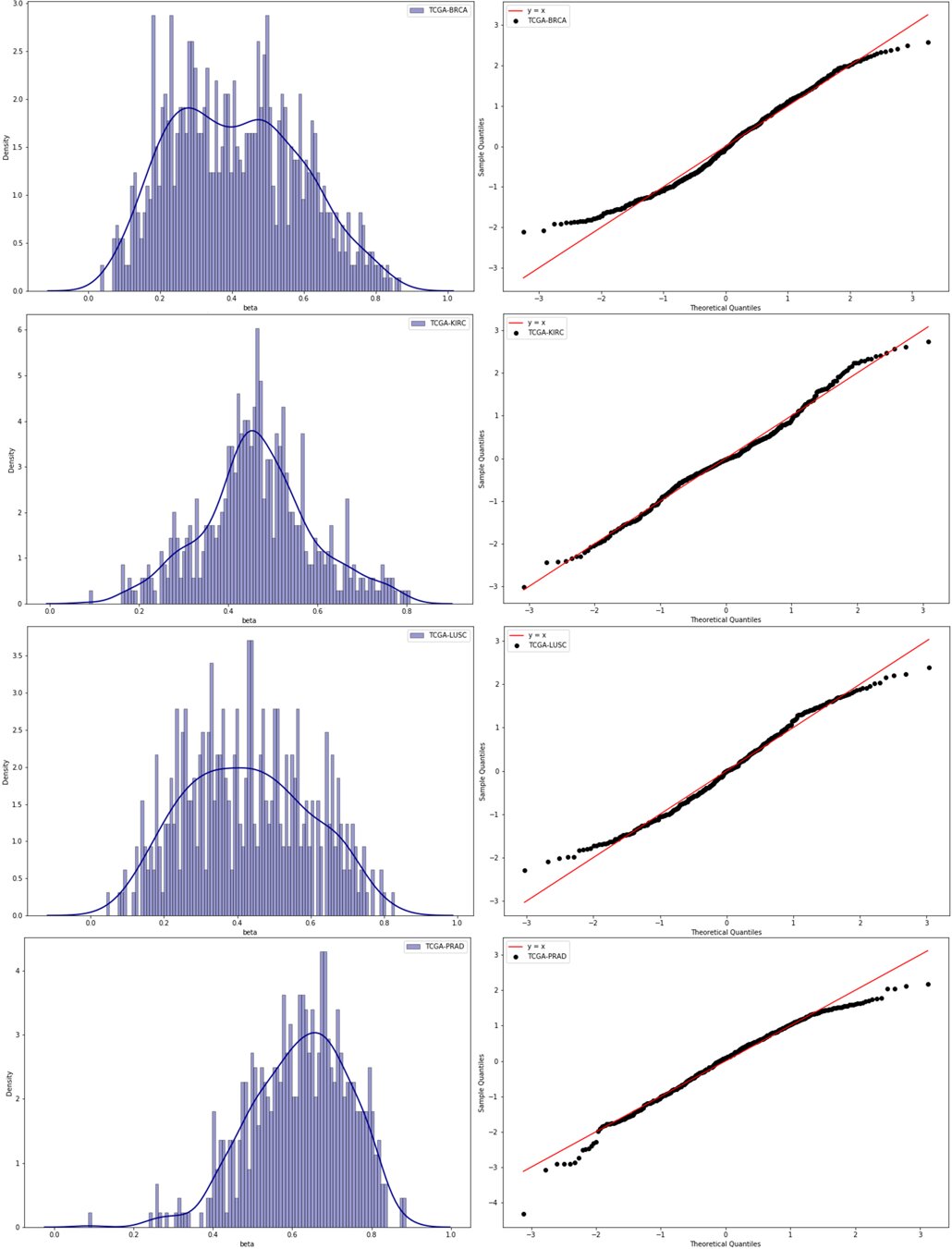

### Supplementary Figure 20

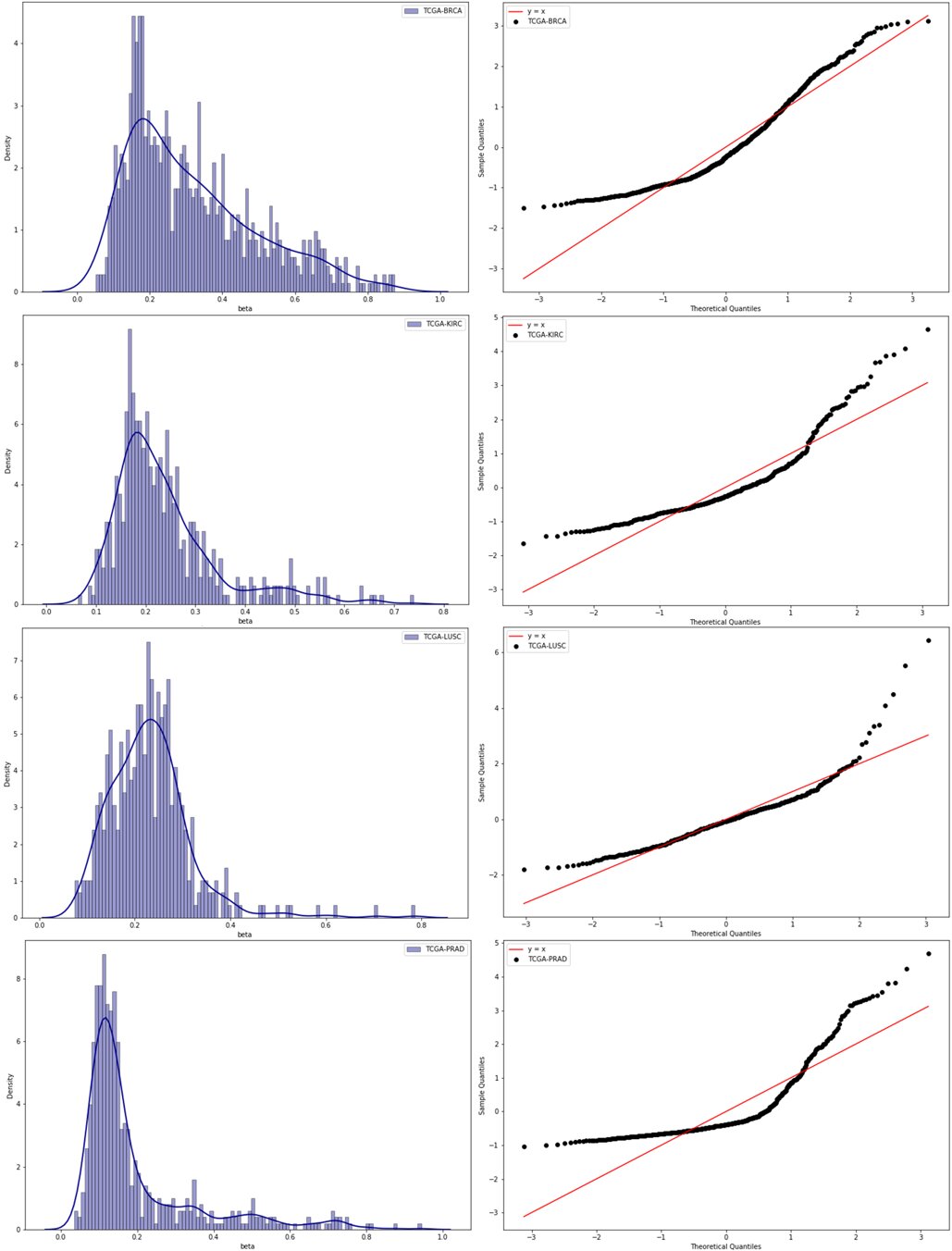

### Supplementary Figure 21

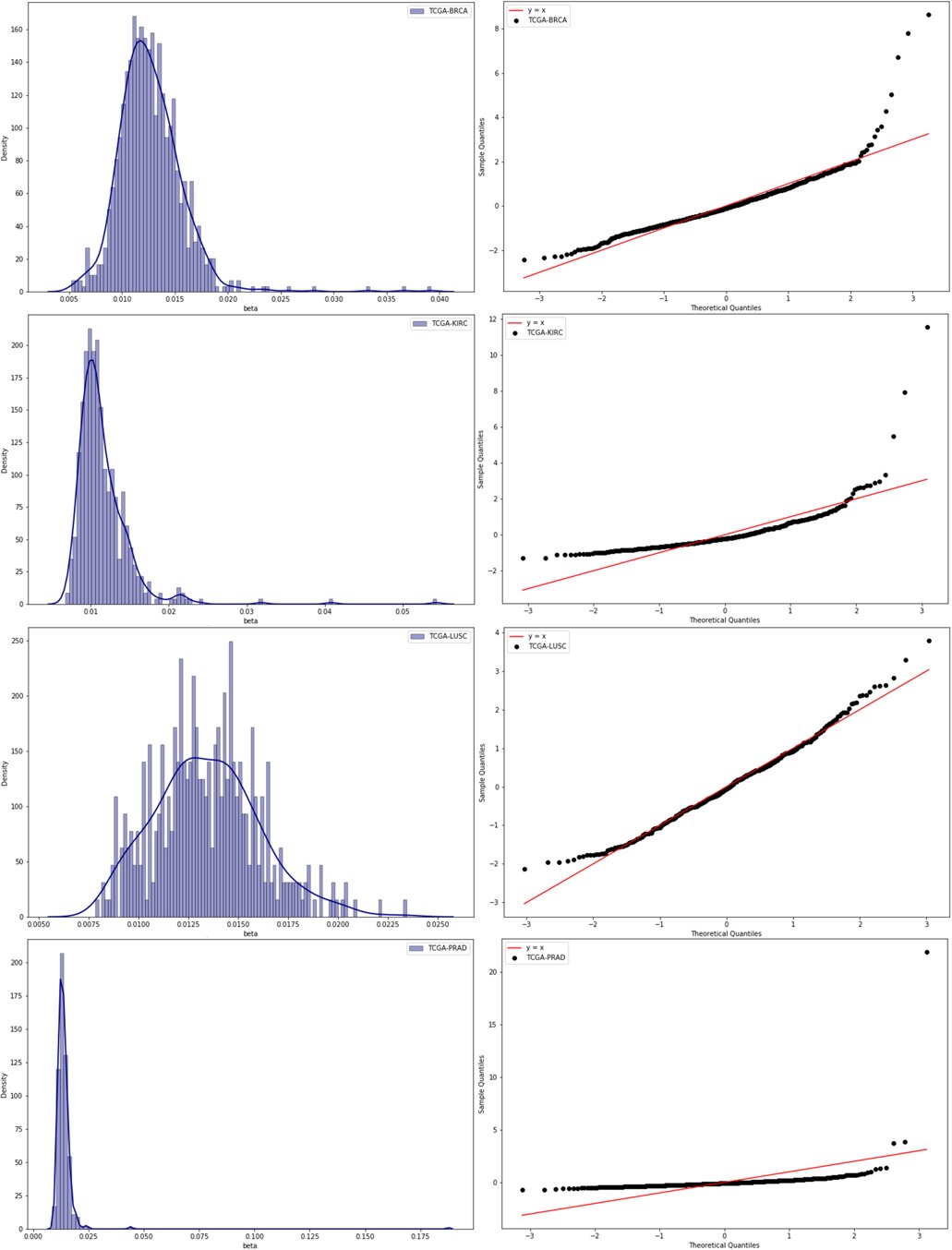

### Supplementary Figure 22

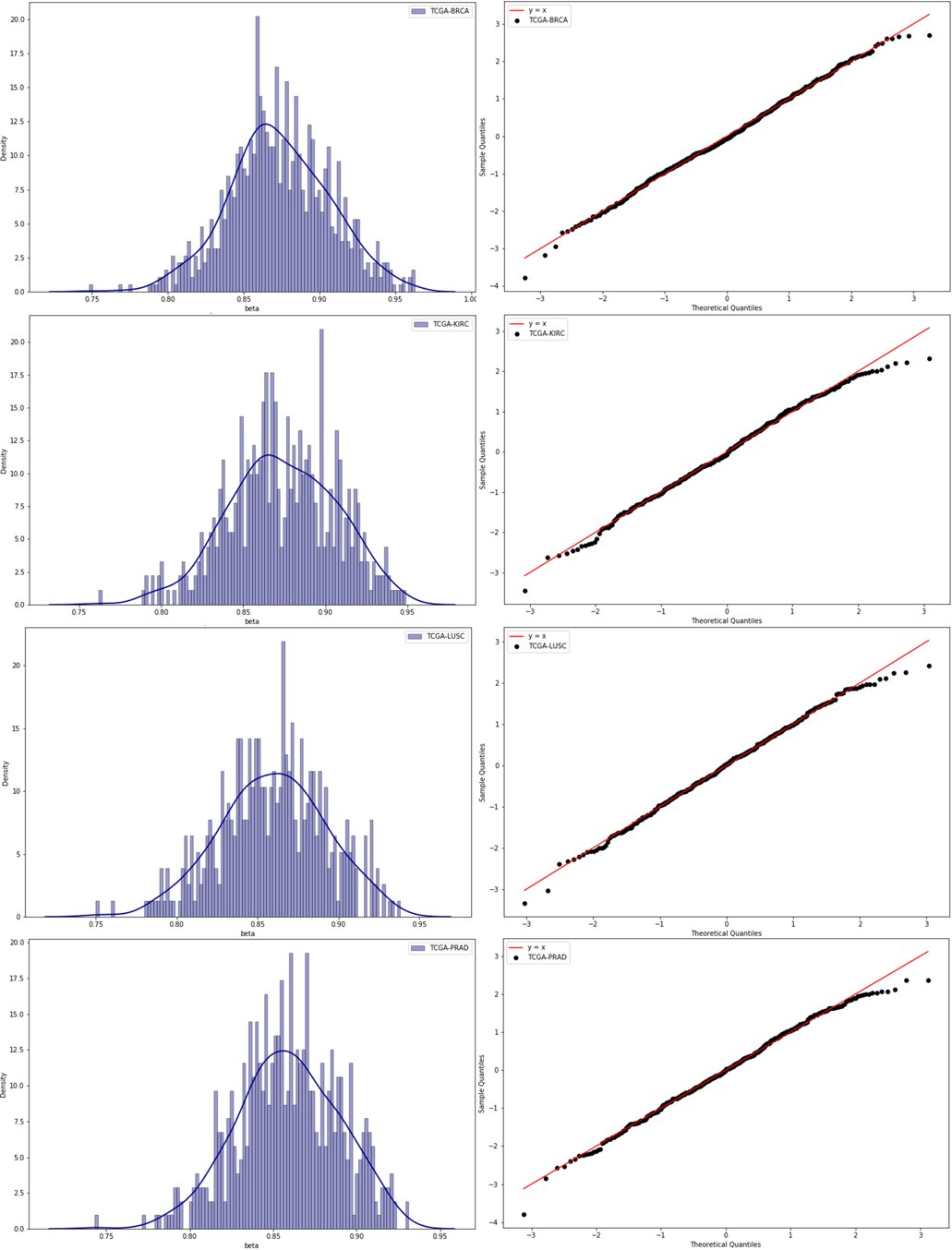

### Supplementary Figure 23

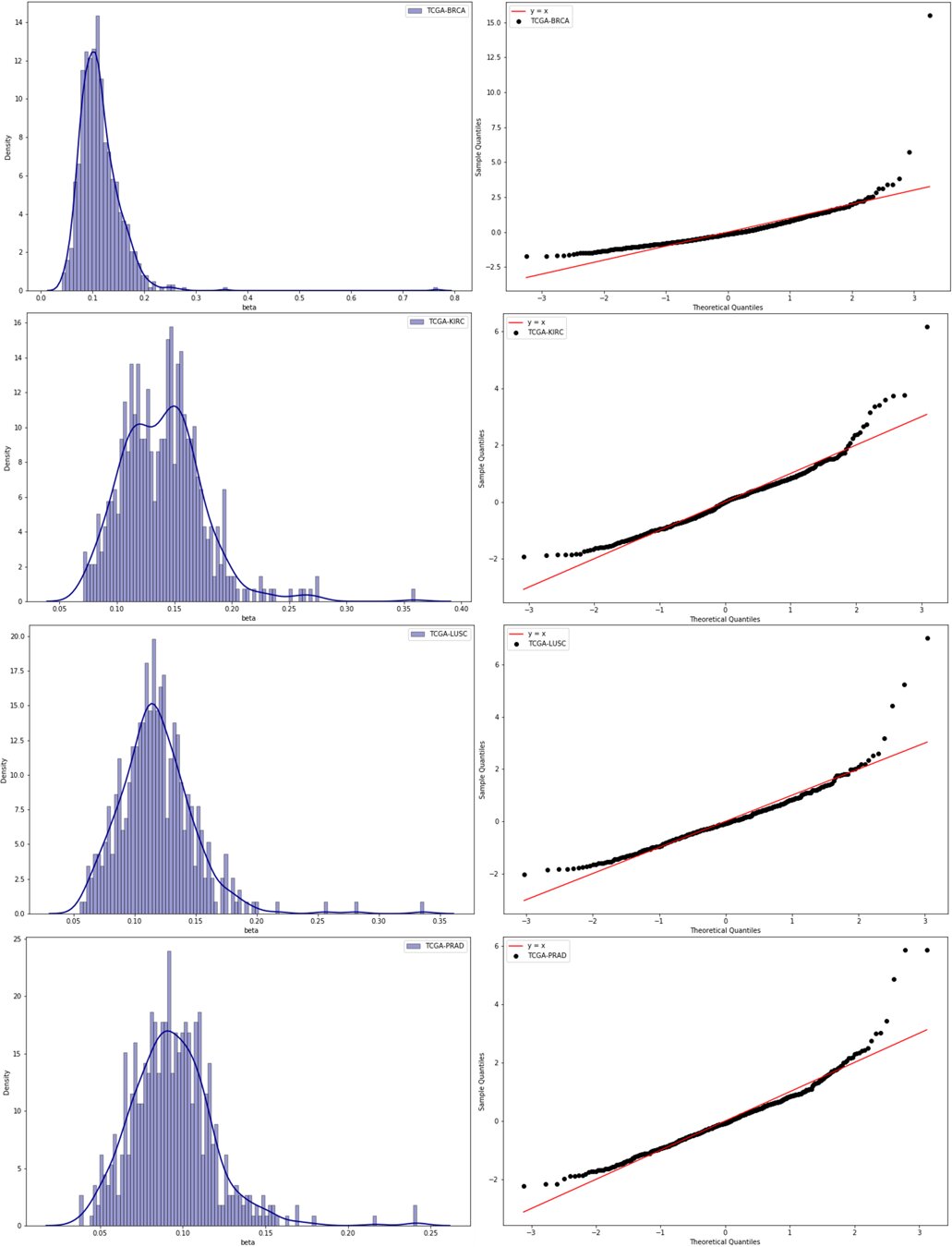

### Supplementary Figure 24

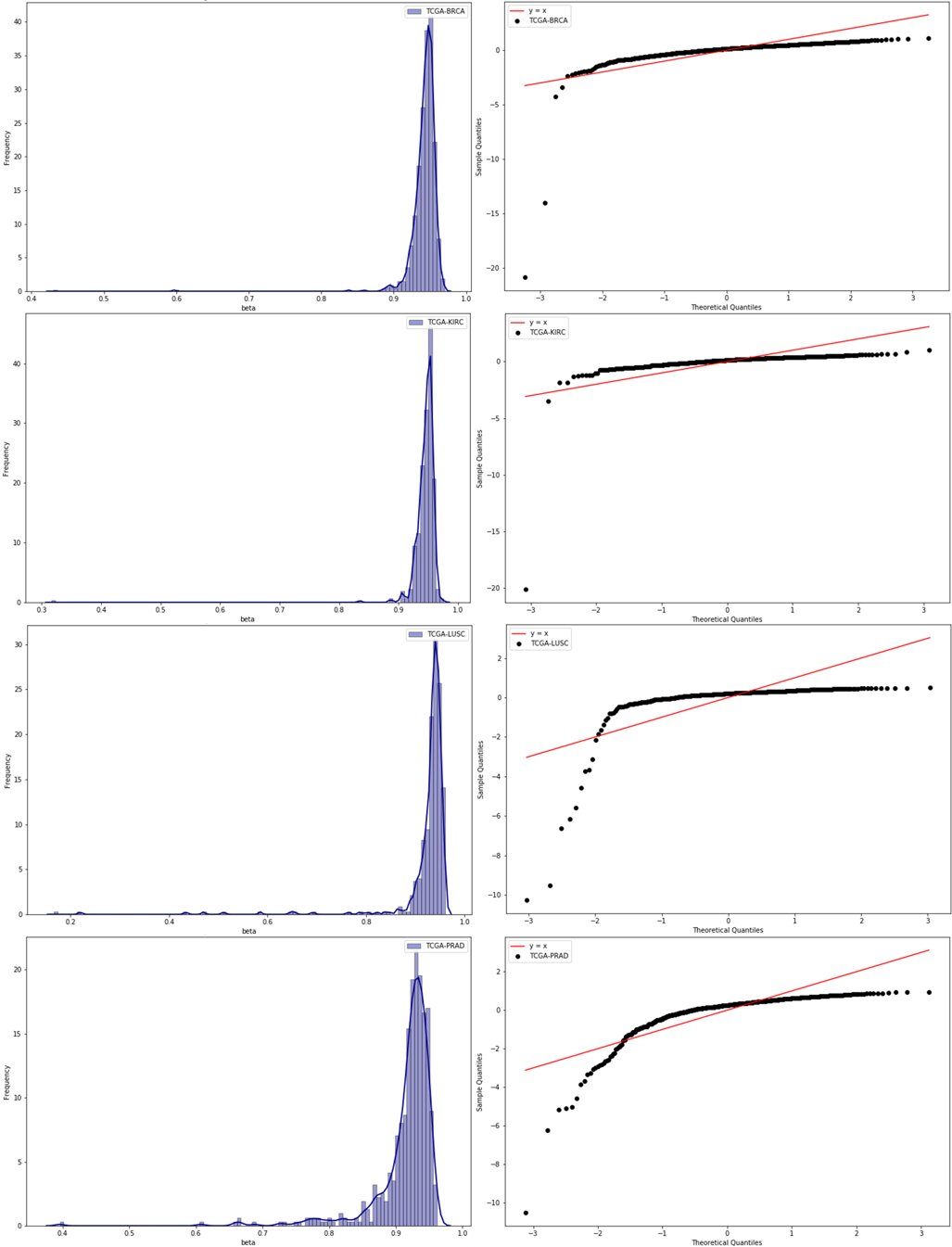

### Supplementary Figure 25

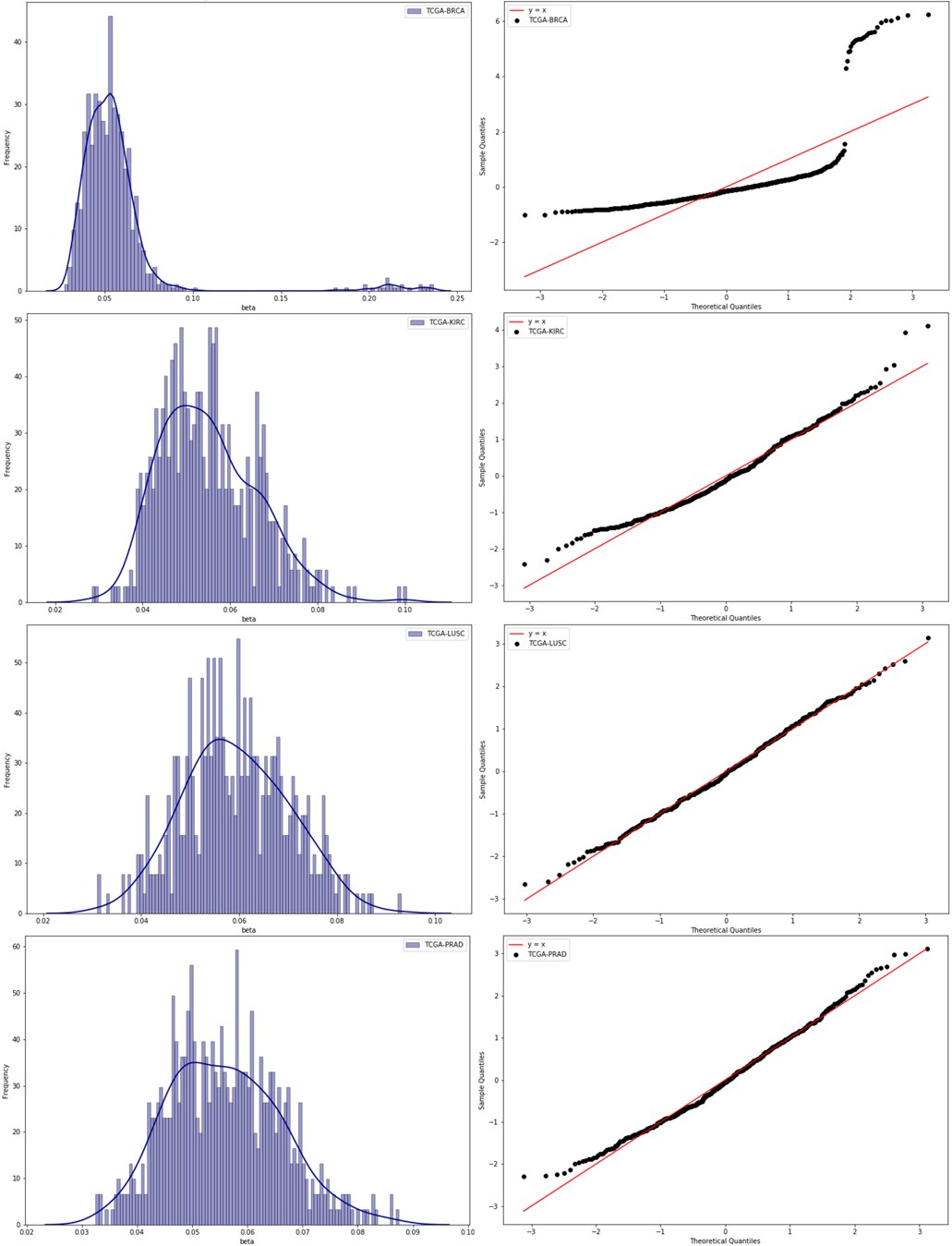

### Supplementary Figure 26

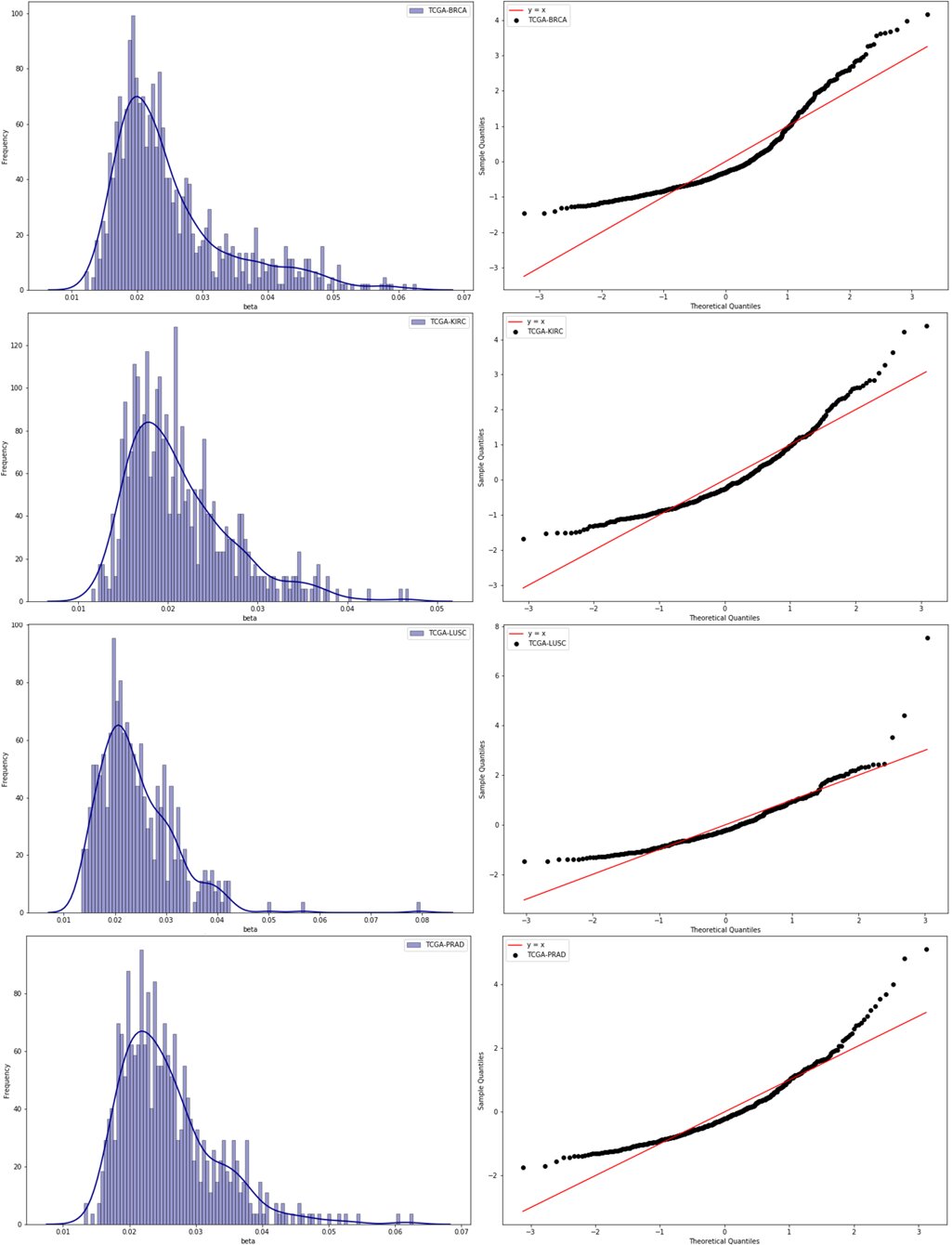

### Supplementary Figure 27

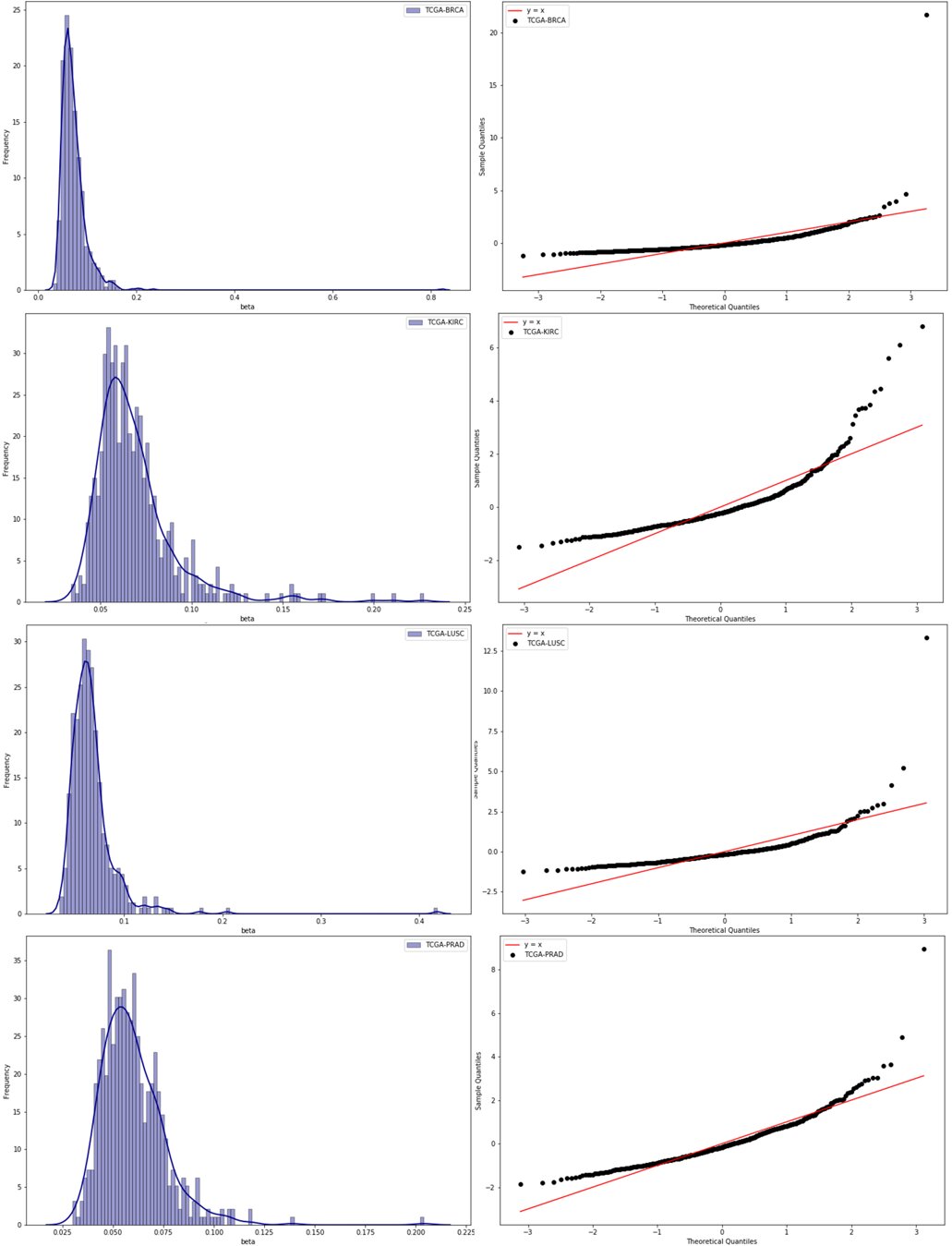

### Supplementary Figure 28

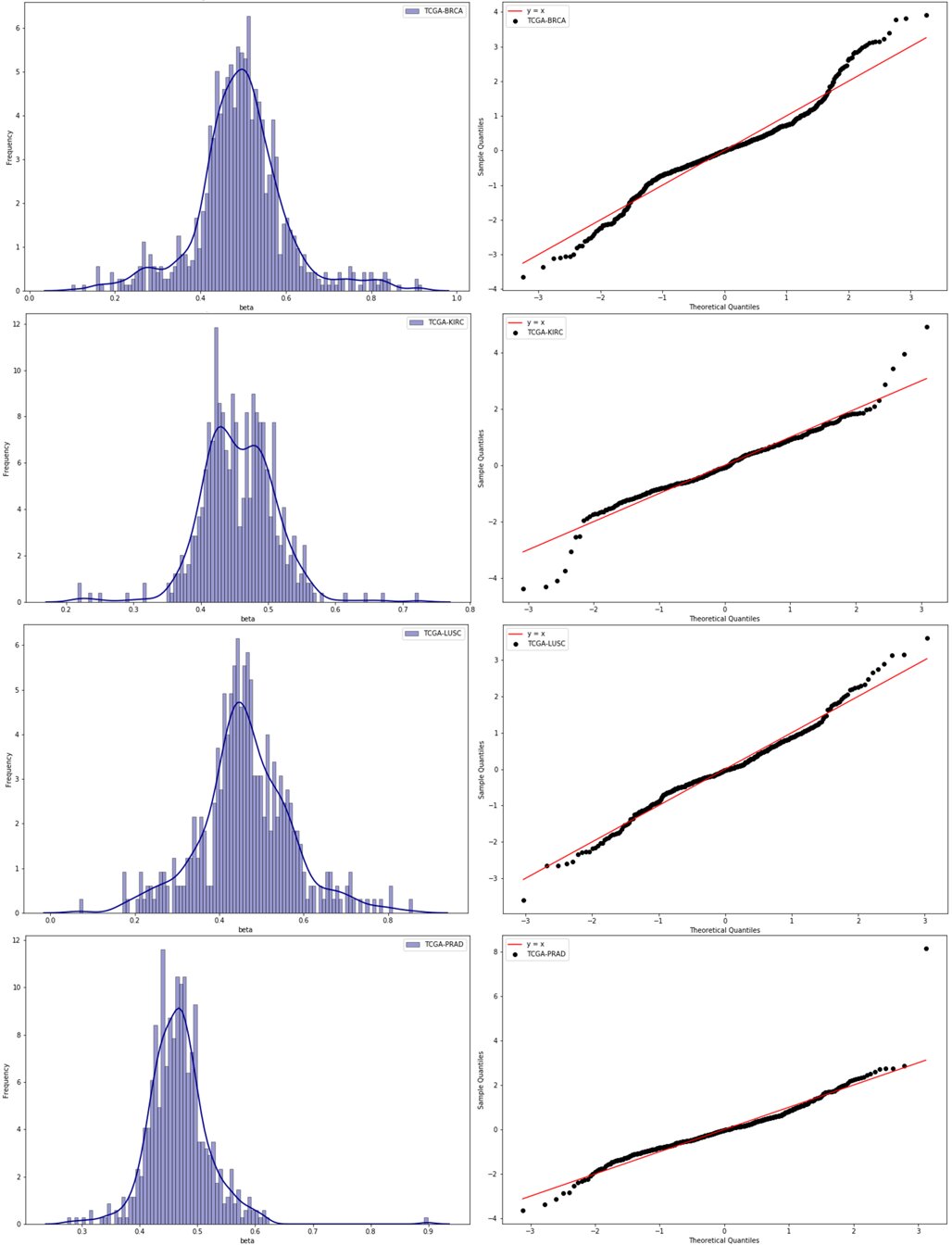

### Supplementary Figure 29

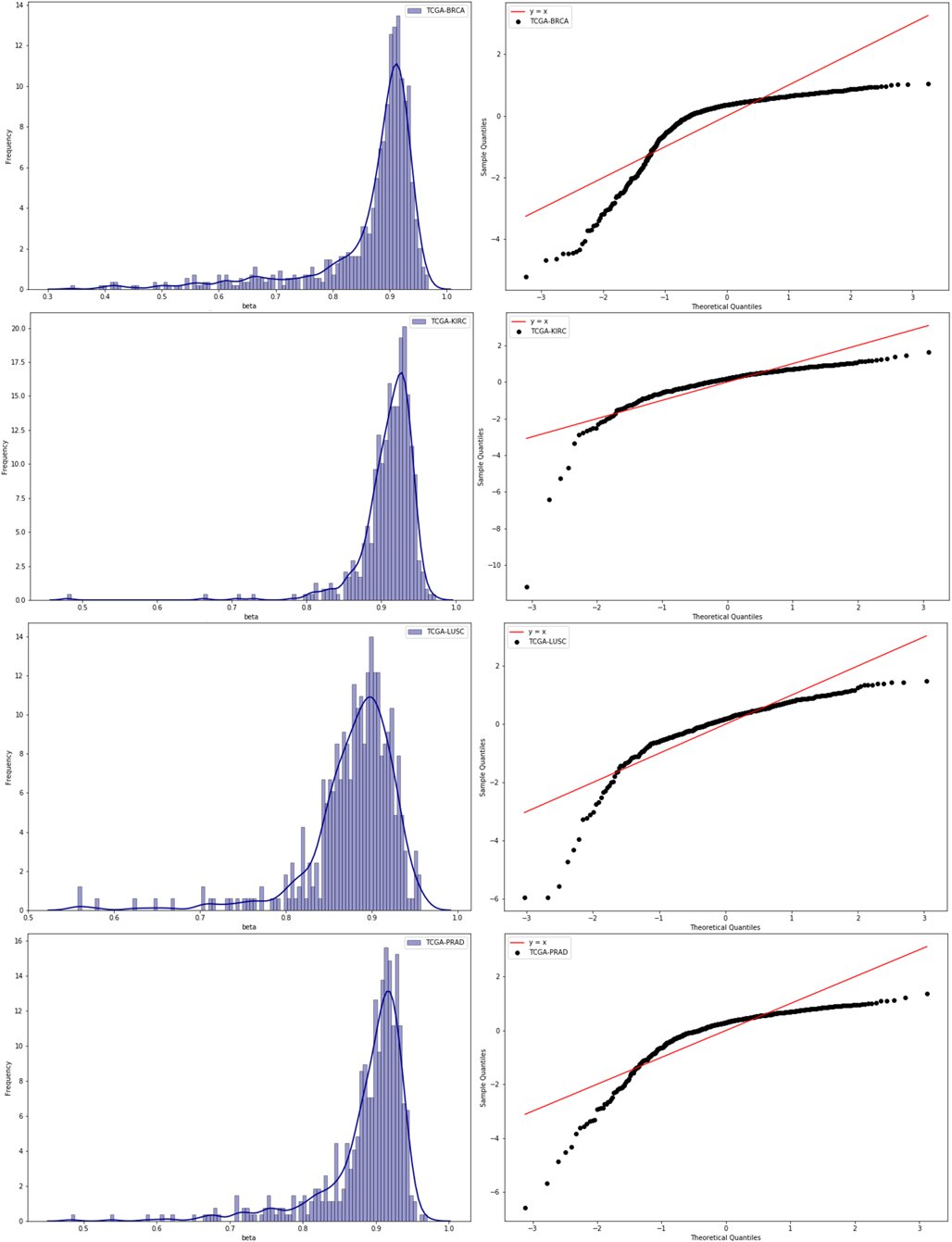

### Supplementary Figure 30

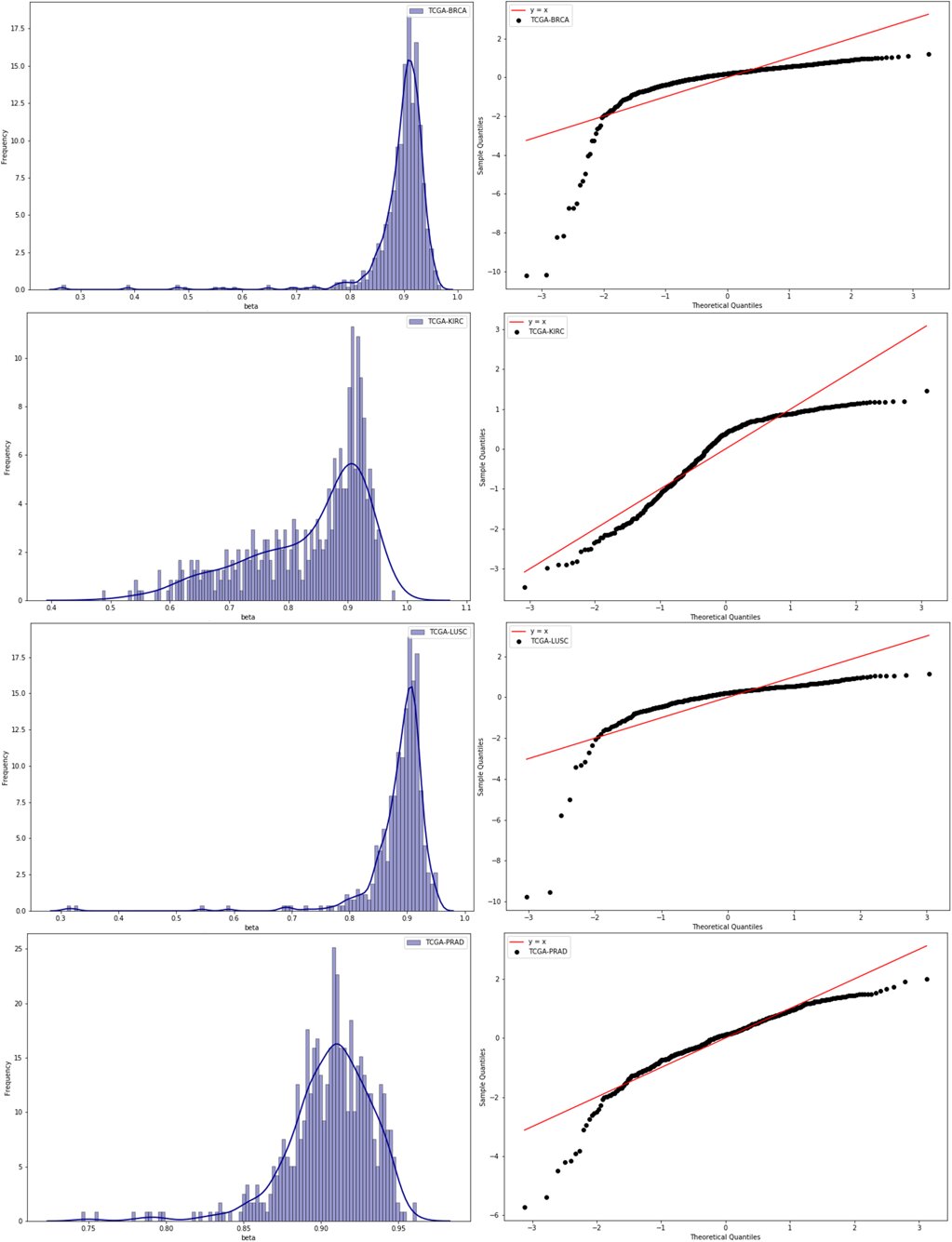

### Supplementary Figure 31

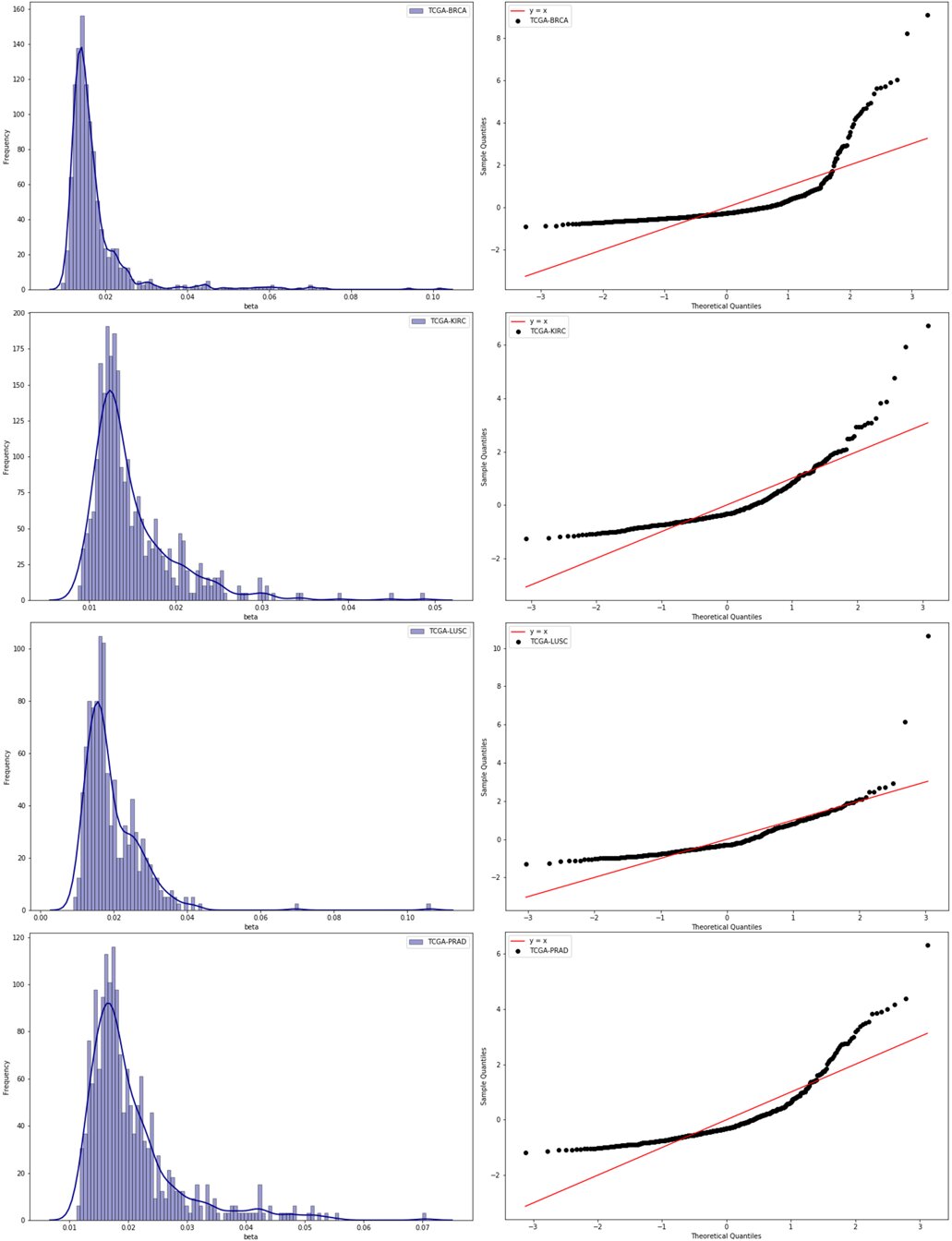

### Supplementary Figure 32

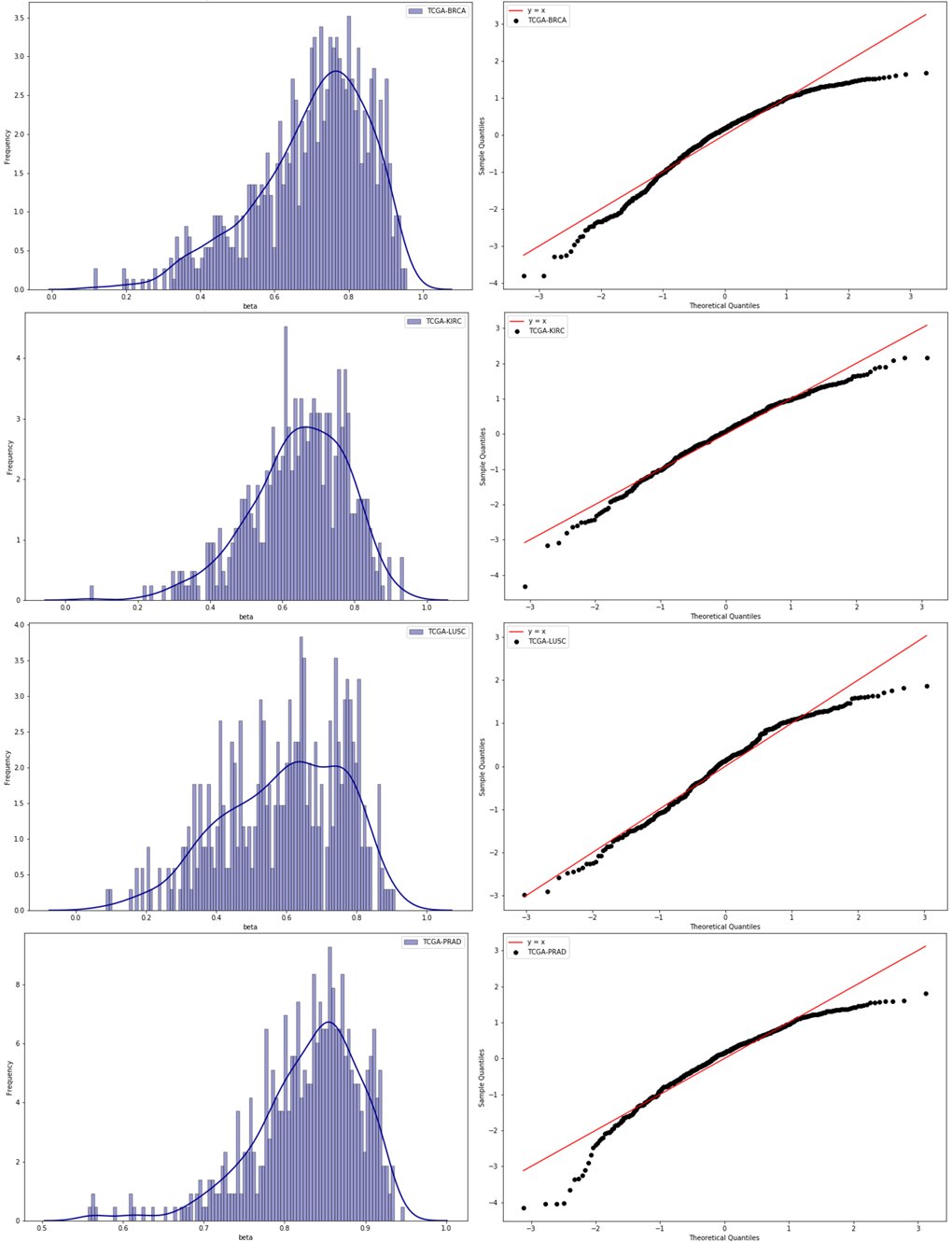

### Supplementary Figure 33

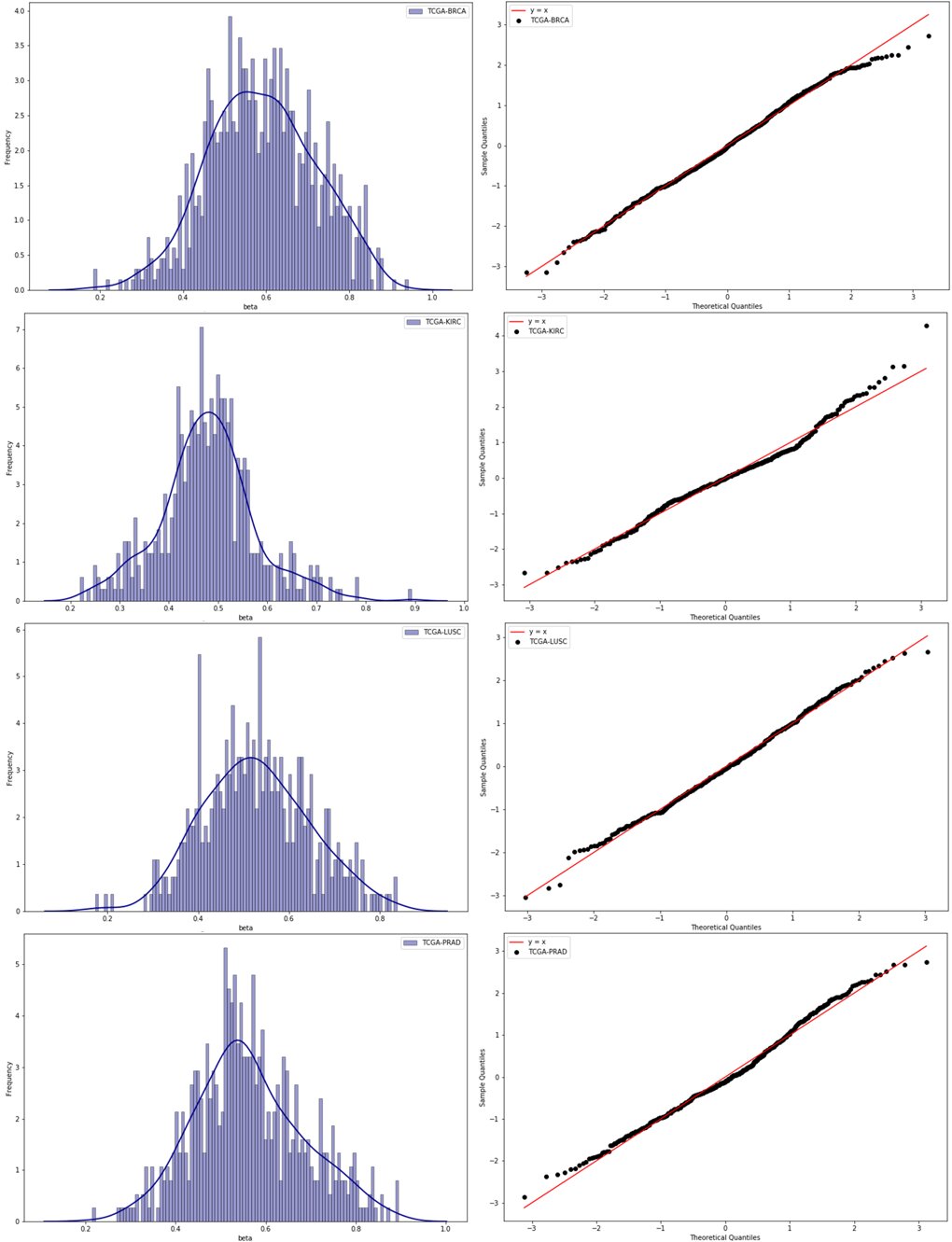

### Supplementary Figure 34

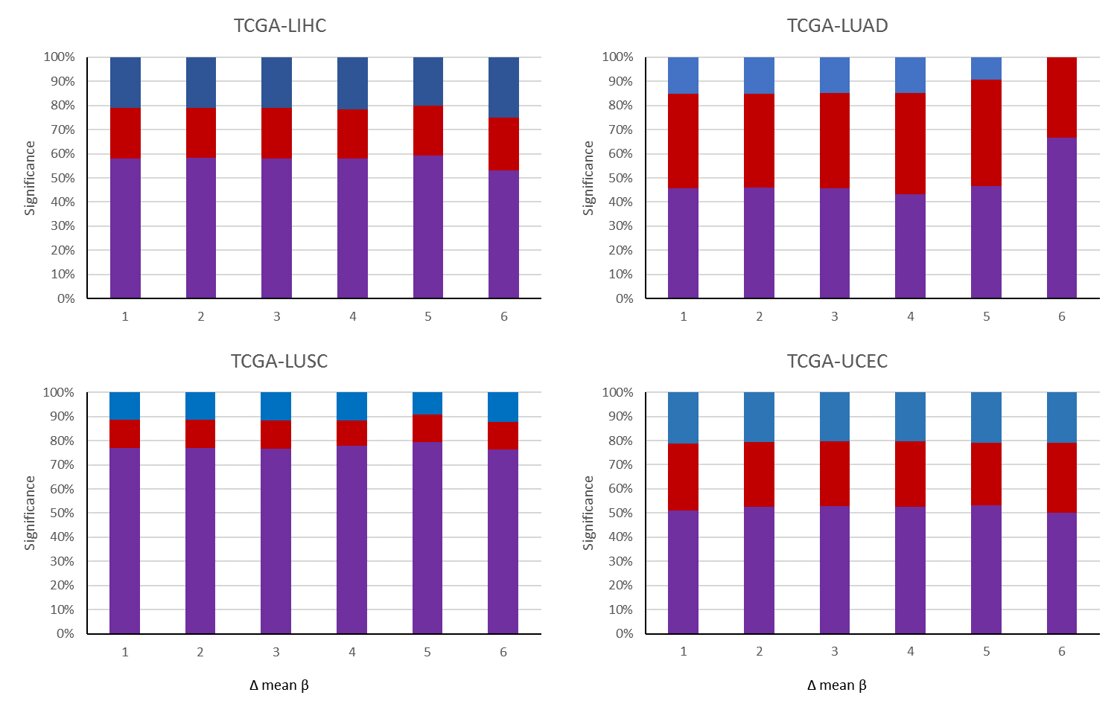
